# Polymorphisms in the yeast galactose sensor underlie a natural continuum of nutrient-decision phenotypes

**DOI:** 10.1101/126011

**Authors:** Kayla B. Lee, Jue Wang, Julius Palme, Renan Escalante-Chong, Bo Hua, Michael Springer

## Abstract

In nature, microbes often need to “decide” which of several available nutrients to utilize, a choice that depends on a cell’s inherent preference and external nutrient levels. While natural environments can have mixtures of different nutrients, phenotypic variation in microbes’ decisions of which nutrient to utilize is poorly studied. Here, we quantified differences in the concentration of glucose and galactose required to induce galactose-responsive (GAL) genes across 36 wild *S. cerevisiae* strains. Using bulk segregant analysis, we found that a locus containing the galactose sensor *GAL3* was associated with differences in GAL signaling in eight different crosses. Using allele replacements, we confirmed that *GAL3* is the major driver of GAL induction variation, and that *GAL3* allelic variation alone can explain as much as 90% of the variation in GAL induction in a cross. The *GAL3* variants we found modulate the diauxic lag, a selectable trait. These results suggest that ecological constraints on the galactose pathway may have led to variation in a single protein, allowing cells to quantitatively tune their response to nutrient changes in the environment.

**Author summary:** In nature, microbes often need to decide which of many potential nutrients to consume. This decision making process is complex, involving both intracellular constraints and the organism’s perception of the environment. To begin to mimic the complexity of natural environments, we grew cells in mixtures of two sugars, glucose and galactose. We find that in mixed environments, the sugar concentration at which cells decides to induce galactose-utilizing (GAL) genes is highly variable in natural isolates of yeast. By analyzing crosses of phenotypically different strains, we identified a locus containing the galactose sensor, a gene that in theory could allow cells to tune their perception of the environment. We confirmed that the galactose sensor can explain upwards of 90% of the variation in the decision to induce GAL genes. Finally, we show that the variation in the galactose sensor can modulate the time required for cells to switch from utilizing glucose to galactose. Our results suggest that signaling pathways can be highly variable across strains and thereby might allow for rapid adaption in fluctuating environments.

## Introduction

The nutrient composition of natural environments can fluctuate and organisms must induce metabolic pathways that allow them to utilize the available nutrients [1-3]. Recent studies have found that closely related microbes vary in both the types of nutrients they can utilize and the efficiency at which they do so [4,5]. However, most studies have focused on differences in growth in single-nutrient environments (e.g., growth in “pure” glycerol). Natural environments, on the other hand, often contain multiple nutrients that cells need to choose between, and suboptimal nutrient decisions can have severe fitness consequences [6-9]. Hence, it is likely that cells have been selected not only to utilize nutrients efficiently, but to decide which subsets of nutrients to utilize.

Signaling pathways sense which nutrients are present and control the decision of which transcriptional network to activate. The majority of plasticity in gene expression patterns has been linked to changes in transcriptional regulatory networks. While transcription factor binding sites are typically conserved, the location of the binding sites in the genome can rapidly evolve [10]. Chromatin immunoprecipitation followed by sequencing in yeast [11-13], mice and human [14], and flies [15] have shown a surprisingly small conservation in the genes and sites that were bound by transcriptional regulators between species. Even when the regulated genes are conserved, the transcription factors that regulate them can change [16-19]. The development of genomic tools has greatly aided the interspecific comparison of regulatory binding sites.

There are relatively few cases where adaptive changes in signaling networks have been linked to molecular and genetic variation. By contrast, changes in transcription regulatory networks have been easier to identify due to the development of high-throughput genomic and computational approaches. Additionally, studies are often biased towards finding changes in transcriptional regulatory networks based on the phenotypes assayed, i.e. fitness in 'extreme' environments. Still, there are multiple situations where upstream signaling changes must have occurred. For instance, in the galactose-utilization pathway (GAL) in *C. albicans*, Rtg1p and Rtg3p activate GAL genes while Gal4p is involved in glucose regulation; in *S. cerevisiae*, Gal4p activates GAL genes, while Rtg1p and Rtg3p are involved in glucose regulation [20]. This implies that the upstream signaling networks that sense and transduce galactose and glucose signals have also changed. Furthermore, duplication and divergence can shape signaling networks. For example in the GAL pathway in yeast, duplication and divergence allowed the sensing and catabolic activity of a single ancestral protein to be separated into two paralogous proteins [21]; this divergence likely had profound consequences for how yeast were able to ‘perceive’ galactose. Hence, it is likely that cellular decision-making can also evolve, but the degree of variation, its molecular and physiological basis, and the evolutionary timeframe at which it occurs has yet to be resolved.

To begin to address these questions we characterized differences in natural isolates of the budding yeast, *S. cerevisiae* in the *decision* to induce the GAL pathway in mixtures of glucose and galactose. In the presence of high concentrations of glucose, the preferred carbon source, yeast cells repress the GAL pathway [22,23]. In the presence of galactose alone, cells activate GAL genes. In mixtures of both glucose and galactose, cells must “decide” whether to induce GAL-associated genes. In such mixed environments, cells show a complex response [24] where the induction of the pathway is dependent on the ratio of glucose and galactose [25]. These observations, combined with the deep molecular understanding in the literature [26,27], make the GAL pathway an excellent model for studying natural variation in cellular decision-making.

Here, we use single-cell measurements to quantify differences in GAL decision-making across closely related natural isolates of *S. cerevisiae*, followed by bulk segregant analysis and allele replacements to find the genetic determinants of this variation. We found that the glucose concentration needed to induce GAL genes varies ∼100-fold across yeast strains. Even though this phenotypic variation is continuous, a large proportion of it can be explained by differences in a single gene, the galactose sensor *GAL3*. Changing the *GAL3* allele produces a measurable difference in the diauxic lag length, a trait that was previously shown to be selectable [8]. These results highlight the fact that cellular decision-making has the potential to be rapidly shaped by selective pressures in the environment.

## Results

### The decision to induce GAL pathway varies across *S. cerevisiae* natural isolates

To enable measurement of the GAL signaling response, we generated a fusion of the *GAL1* promoter from *S. cerevisiae* and yellow fluorescent protein (*GAL1pr-YFP*) (Fig. 1A). *GAL1* is the first metabolic gene in the galactose utilization pathway [28] and this promoter has been used by numerous studies as a faithful readout of pathway activity [9,25,29,30]. The reporter construct was integrated into the neutral HO locus [31] in 42 different *S. cerevisiae* strain backgrounds (S1 Fig.) [32,33]. These 42 strains span a range of phylogenetic and ecological diversity [32,33]. Six of these strains either did not grow in galactose, likely due to inactivation of the pathway [4], and thus were not characterized further. We focused on determining the GAL response phenotype of the remaining 36 strains (S1 Table).

**Figure 1.**
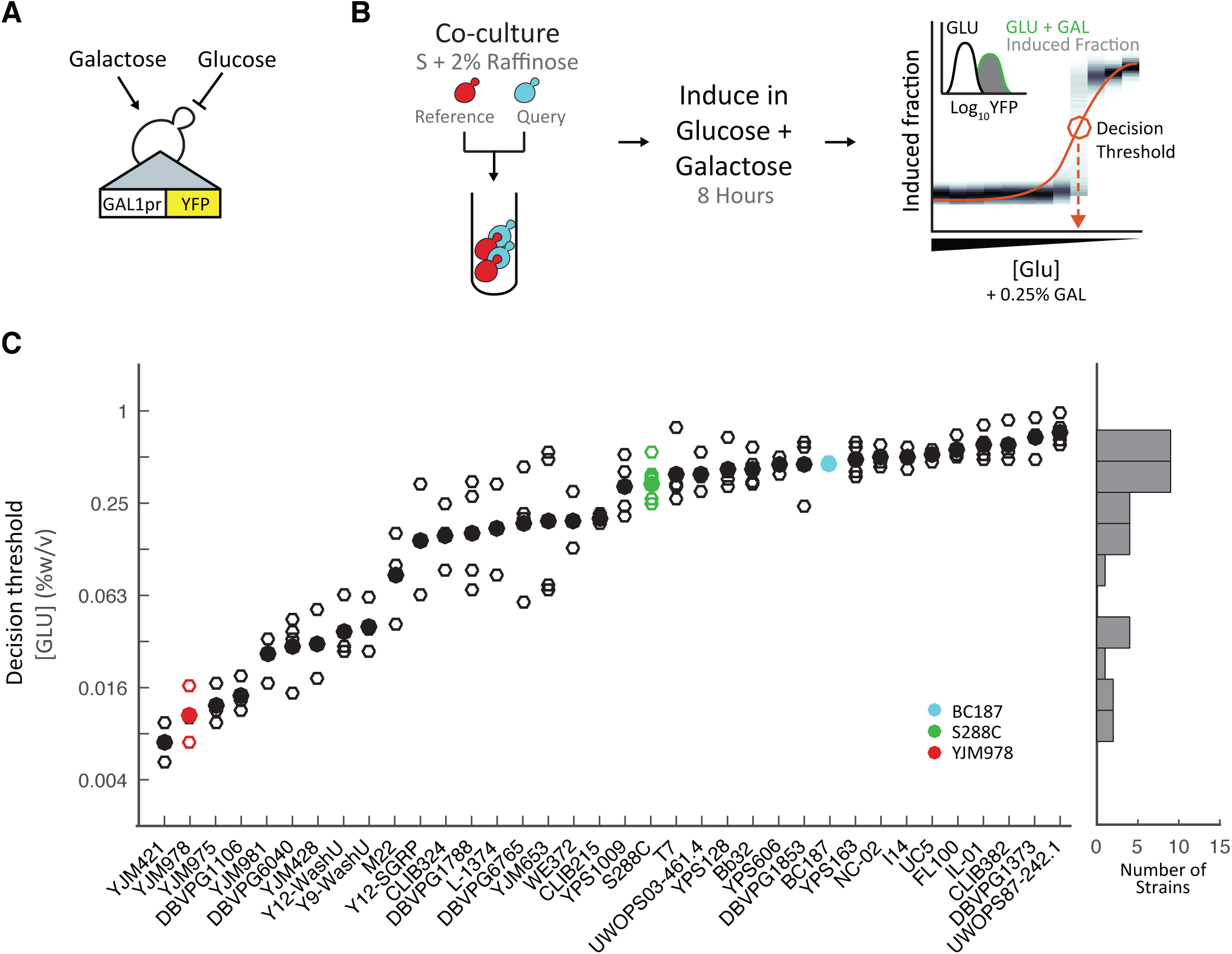
Natural isolates of *S. cerevisiae* vary in the decision to induce the GAL pathway. (A) Schematic of the reporter construct (*GAL1pr-YFP*). (B) Schematic of co-culture pre-growth, glucose and galactose induction, and flow cytometry measurement for a glucose titration. The decision threshold, the concentration of glucose where 50% of the cells are induced, is indicated by circle and dashed line. (C) Decision threshold for 36 lab and natural isolates of *S. cerevisiae*. Histogram shows the distribution of the mean decision threshold for all strains assayed.

To survey the natural variation in the inducibility of GAL genes in mixtures of glucose and galactose, we measured the GAL reporter response in a titration of glucose concentrations from 2% to 0.004% w/v on a background of constant 0.25% galactose (Fig. 1B, Materials and methods). Cells were first pre-grown for 14-16 hours in 2% raffinose (which does not induce or repress the GAL pathway), and then transferred to glucose + galactose and grown for 8 hours at low densities. We previously showed that this protocol is sufficient for cells to reach steady-state without depleting the carbon sources [25]. Finally, single-cell YFP fluorescence was measured by flow cytometry. To account for well-to-well variability or variability in our glucose titration, each of the 36 query strains were co-cultured with a reference strain, YJM978, containing *TDH3pr-mCherry* (this constitutive fluorophore allowed us to distinguish the query and reference strains) and *GAL1pr-YFP* (Materials and Methods).

Qualitatively, there were large strain-to-strain differences in the concentration of glucose at which cells induced the GAL pathway (Fig. 1, S4 Fig.). We also observed bimodal expression in some strains and conditions, a likely consequence of cellular heterogeneity and ultrasensitivity in the GAL circuit [30,34,35]. This complicates quantitative analysis, because a metric such as the mean expression (which is implicit, for example, in a bulk assay) would convolute both the number of cells that are inducing and the expression level of the cells that have ‘decided to’ induce, two factors that may vary independently in bimodal responses. Hence, to compare the GAL pathway response between natural isolates, we defined a metric, the “decision threshold”, as the concentration at which 50% of cells have greater-than-basal expression of the GAL reporter (Materials and Methods). This metric is similar to those used in previous work [29,36], and focuses on *when* a cell decides to induce a pathway while differentiating it from *how strongly* a cell responds once induced. The decision threshold is highly reproducible across replicate measurements for all of our natural isolates (S3 Fig.).

Quantitatively, the decision threshold varies over a range of 108 ± 0.7-fold glucose concentrations across our strains (Fig. 1, S4 Fig.). The Hawaiian cactus strain UWOPS87-242.1, was most inducible, with a decision threshold of 0.74±0.2% glucose (mean ± standard error mean), while the clinical isolate, YJM421, was least inducible, with a decision threshold of 0.01±0.01% glucose (mean ± S.E.M.). Half of the strains have decision thresholds within a 8.1- fold range centered at 0.25% glucose (Fig. 1C). This glucose concentration corresponds to a galactose+glucose ratio of ∼1:1. The distribution of decision thresholds appears continuous; there are significantly more than two distinct decision thresholds given the reproducibility of our measurements (Materials and Methods).

Strain differences in decision threshold could be due to differences in sugar signaling, utilization, or both. If sugar utilization is a factor, we expect the decision threshold to be correlated to growth rates in glucose or galactose. We measured the growth rates of the 36 natural isolates during mid-exponential growth in either 0.5% glucose or 0.5% galactose (S5 Fig., [6]). Despite substantial variation in single-sugar growth rates across our strains (0.23-fold in glucose and 0.16-fold in galactose), neither growth in pure galactose or glucose are correlated with the decision threshold (glucose r^2^ = 0.2, galactose r^2^ = 0.001). This implies that while both sugar utilization and signaling can vary between strains, evolution has the potential to select these two traits independently.

Previous studies have determined the correlation between genotypic diversity and either phenotypic diversity or ecological niche. For example, analysis of 600 traits in yeast by Warringer et al. identified a correlation between phylogeny and phenotype [4]. These studies can be used to assess whether traits are more likely to be neutral or undergoing selective constraint. To determine if the decision threshold is correlated with phylogeny, we began by comparing the 13 closely related strains of the wine/European clean lineage. Despite the close phylogenetic relationship of these strains, this lineage represents the most phenotypic diversity. The two most phenotypically distinct strains in this lineage, YJM978 and DBVPG1373, have a 48±0.3-fold difference (mean ± S.E.M.). More broadly, we compared the pairwise genetic distances (determined by RAD-SEQ [37]) to pairwise phenotypic distance (Materials and Methods). We did not find a significant correlation (r^2^=-0.08) between genetic distance and decision threshold. This level of correlation with genetic distance is comparable to that of many other traits [4] (S6 Fig., p-value=0.17 by ANOVA). Finally, we tested for and found a signification association between ecological niche and decision threshold (S7 Fig., p-value=2.95e-5 by ANOVA).

### Bulk segregant analysis identifies one major-effect locus underlying natural variation in the GAL decision threshold

To investigate the genetic basis of the observed variation in GAL decision threshold, we performed bulk-segregant analysis using a variant of the X-QTL method (Fig. 2A) [38-40]. We crossed eight strains that span the phenotypic and phylogenetic diversity of *S. cerevisiae* in a round-robin design (Fig. 2). This design is known to efficiently sample parental genetic variation and allow downstream linkage analyses to detect loci with a range of effect sizes [38]. Pools of segregants from each cross were grown in a glucose + galactose condition that maximally differentiates the parental phenotypes. The 5% least and 5% most induced cells (“OFF” and “ON” segregant pools) were isolated by fluorescence-activated cell sorting (FACS) and sequenced in bulk to determine the parental allele frequencies in each pool. We used the MULTIPOOL software [41] to determine statistical significance for allele frequency differences between OFF and ON pools across the genome (Materials and Methods), and called significant loci as regions where the peak log-odds-ratio was greater than 10 (Fig. 2B). This cutoff had a low false-discovery rate in a previous study, and correlated well with allele frequency difference, a proxy for locus effect size, in our data [38] (S8 Fig.).

**Figure 2.**
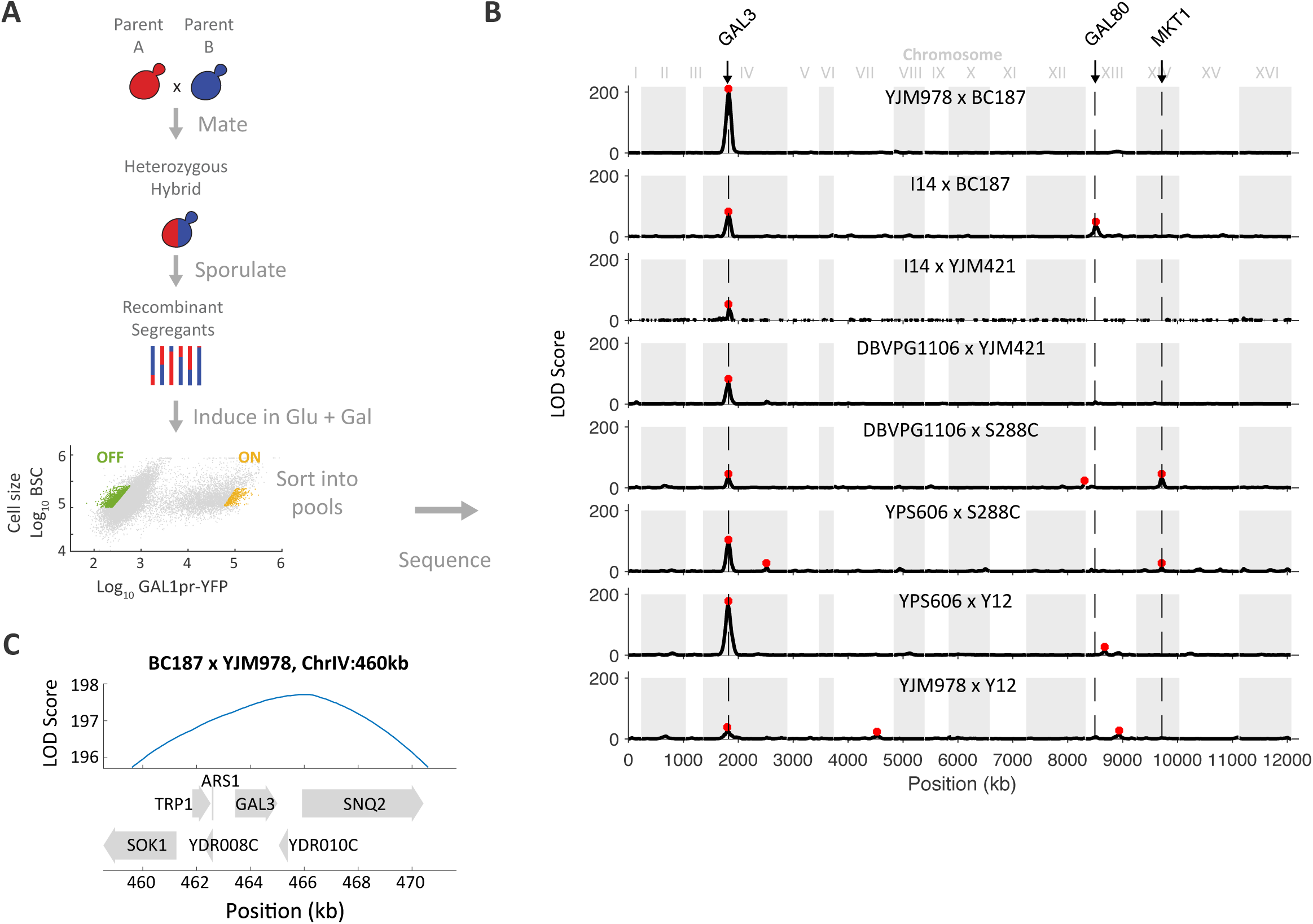
Bulk segregant analysis identifies one major-effect locus underlying natural variation in decision threshold. (A) Schematic of bulk segregant analysis. Meiotic segregants from heterozygous hybrids were sorted by FACS into ‘ON’ and ‘OFF’ pools based on *GAL1pr-YFP* expression and then sequenced. (B) LOD score of allele frequency difference between ‘ON’ and ‘OFF’ segregant pools versus genomic position (red asterisks: LOD > 10). A region of chromosome IV containing *GAL3* was associated with the difference between the ‘ON’ and ‘OFF’ phenotype in all 8 crosses. Potential candidate genes for other loci include *GAL80, MKT1*, and others listed in Table S2. (C) Genes found within the 2-LOD support interval around the peak LOD score from ChrIV:460kb plotted with the LOD score for the BC187xYJM978 cross.

Over all 8 crosses, we found 16 loci where segregant pools differ in allele frequency at LOD > 10 (Fig. 2B). One locus centered at 460 kb on chromosome IV (henceforth, “chrIV:460”) was the only locus to exceed the LOD cutoff in all 8 crosses, as well as the most significant locus in each cross (Fig. 2B). The 2-LOD support interval for this locus in the YJM978 x BC187 cross, defined as the genomic region where LOD decreases by 2 from its peak, is 10 kb wide and contains six genes (Fig. 2C). This includes *GAL3*, whose product directly binds galactose and positively regulates the GAL pathway [42]. The support interval for chr:460 looked similar in other crosses (S2 Table). One other locus, at chrXIV:462, reached LOD > 10 in two crosses; the remaining significant loci were confined to a single cross. We did detect additional loci in multiple crosses using a less stringent cutoff of LOD > 5; however, chrIV:460 remained the only locus significant in all crosses (S8 Fig., S2 Table, Materials and Methods).

In principle, a round-robin cross design is expected to detect each locus in more than one cross. The fact that we identified several alleles in only one cross is potentially explained by a lack of statistical power, epistasis, or gene-by-environment effects [38]. Indeed, a potential caveat to the apparent importance of *GAL3* is that in a pooled segregant analysis, large effect QTLs might mask the presence of smaller-effect genes [43]. Furthermore, low sequencing depth of some of our segregant pools may have limited our power to detect small-effect alleles (Materials and Methods). However, even the lowest sequencing depth we obtained (25x) is still sufficiently powered to map alleles with effects as low as 5% of phenotypic variance [44]. More importantly, we performed a complementary analysis of segregants to directly measure the contribution of *GAL3* to the phenotypic variance in a cross (see below). Finally, alleles that were identified in only one cross may arise from the different conditions used for sorting each cross (Materials and Methods), i.e. gene-by-environment effects. Given its importance, we chose to focus on the chrIV:460 locus for further characterization.

### *GAL3* is the causative allele and major driver of variation in the GAL signaling response

To determine if *GAL3* was the causative allele on chrIV:460 with a predictable and quantitative impact on the decision threshold, we replaced the endogenous *GAL3* allele of strains YJM978, BC187, and S288C with alleles from eleven natural isolates spanning the observed range of phenotypic variation (Fig. 1). Allele replacements were constructed by deleting the 3283bp *GAL3* locus, which includes 890 bp upstream, 911 bp downstream, and the 1563 bp *GAL3* ORF in haploid parental strains and then replacing the deleted locus with the homologous ∼3283bp *GAL3* locus from other strains using the CRISPR-Cas9 system [45] (Materials and Methods). Replacement of *GAL3* alleles in the YJM978 background recapitulated the ∼95-fold range of decision threshold of the natural isolates that served as *GAL3* allele donors. Additionally, the decision thresholds of allele-replacement and *GAL3* donor strains were well-correlated in this background (r^2^ of 0.58). Similarly, *GAL3* alleles in the S288C background had a ∼55-fold range and r^2^ of 0.60; *GAL3* alleles in the BC187 background had a ∼138-fold range and r^2^ of 0.63. In total, this confirms the significant impact that the *GAL3* locus has on variation in the decision threshold (Fig. 1, Fig. 3A-C, S9 Fig.).

**Figure 3.**
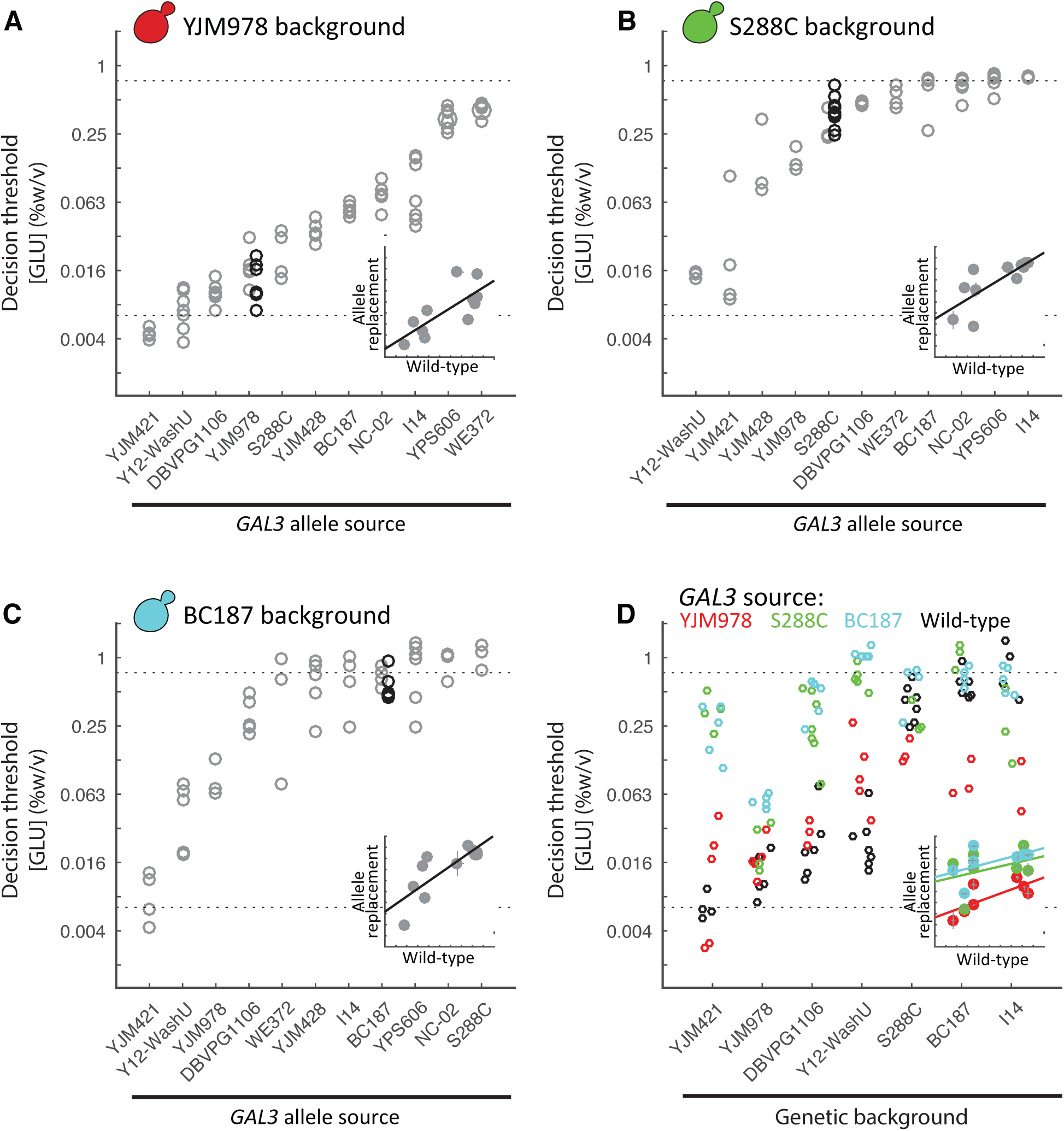
*GAL3* allele largely sets the decision threshold. Decision threshold of eleven different *GAL3* homologous replacements in three genetic backgrounds: (A) YJM978, (B) S288C, and (C) BC187. Decision threshold of wild-type strain is indicated by the black circle. Inset: Correlation plot of natural isolate versus allele replacement decision threshold; error bar represents S.E.M. (D) Decision threshold of three allelic variants of *GAL3* (Mean and S.E.M. of at least two replicates) inserted into various genetic backgrounds: *GAL3^YJM978^* (red), *GAL3^S288C^* (green), and *GAL3^BC187^* (blue), haploid wild-type strain (black). Strains are ordered based on wild-type decision threshold. Inset: Correlation plot of natural isolate (decision threshold of background strain) versus the decision threshold of the allele replacement, error bar represents S.E.M.

While different *GAL3* alleles were able to confer a range of phenotypes in a particular strain background, the three strain backgrounds also displayed different decision thresholds for a given *GAL3* allele. This suggests that genes other than *GAL3* also affect the decision threshold, even for the BC187xYJM978 cross. To assess the magnitude of this background effect, we measured the decision threshold in seven different strain backgrounds where the *GAL3* locus has been replaced with an allele from YJM978, S288C, or BC187 (Fig. 3D, S10 Fig.). Across the seven backgrounds, *GAL3^YJM978^* allele-replacement strains varied in decision threshold over a ∼14-fold range, *GAL3^S288C^* strains over ∼20-fold, and *GAL3^BC187^* strains over ∼49-fold (Fig. 3), and the correlation (r^2^) in decision threshold between these allele-replacement strains and their strain background donors was 0.60, 0.28, and 0.12, respectively. These results confirm that strain background strongly influences decision threshold. However, it is also clear that *GAL3* allele still has a stronger effect, because both the phenotypic range and correlations to donor strain were lower for strain background than for *GAL3* allele. This can also be seen by the fact that the *GAL3^BC187^* and *GAL3^S288C^* strains have decision thresholds that are similar to each other but systematically higher than *GAL3^YJM978^*, regardless of strain background.

### The *GAL3* allele accounts for 70-90% of the phenotypic variance in a cross between strains with extreme opposite decision thresholds

The allele replacements show that *GAL3* is a major driver of natural variation in the decision threshold, but also suggests that other genes play a significant role. To quantify the relative contribution of *GAL3* allele versus other genes to variation in decision threshold, we analyzed the variance in decision threshold across meiotic segregants from YJM978 × BC187 hybrids with different combinations of *GAL3* alleles. This method is relatively insensitive to the metric chosen and potential non-linear relationships between genotype and phenotype. We chose this cross because the *GAL3* locus was the only significant locus from BSA, and thus our calculation should yield a rough upper bound on the *GAL3* contribution in other strains. We constructed three hybrid strains: a ‘wild-type’ hybrid (YJM978 × BC187), a hybrid with *GAL3* only from YJM978 (YJM978 × BC187 *gal3Δ::GAL3^YJM978^*) and a hybrid with *GAL3* only from BC187 (YJM978 *gal3Δ::GAL3^BC187^* x BC187). The decision threshold of at least 58 meiotic segregants was measured for each hybrid in duplicate (Fig. 4, S11 Fig.). Consistent with *GAL3* having a large effect, we found that converting a single allele in each hybrid greatly reduced the phenotypic variation of the segregant populations.

**Figure 4.**
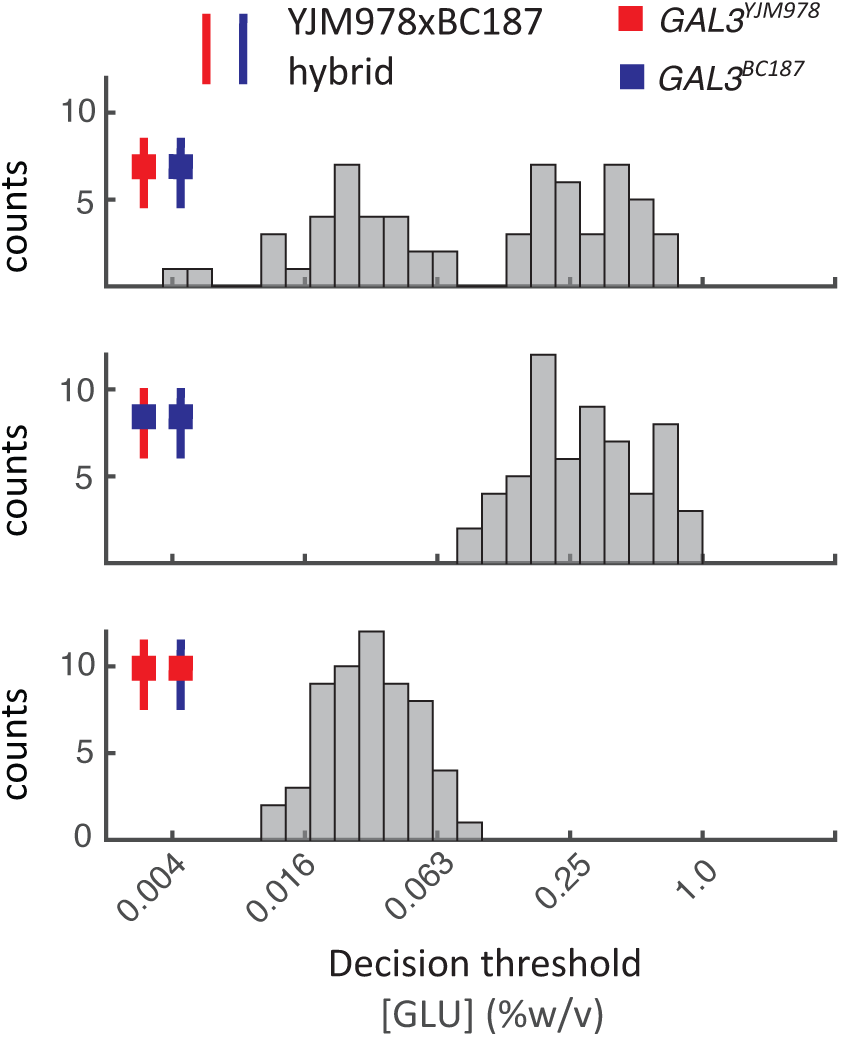
The *GAL3* allele accounts for 70-90% of the decision threshold. Decision threshold of segregants produced from hybrid (top), hybrid with *GAL3^YJM978^* allele homologously replaced with *GAL3^BC187^* (middle), and hybrid with *GAL3^BC187^* allele homologously replaced with *GAL3^YJM978^* (bottom). Hybrids are indicated by small schematic, the line represents the genetic background and the filled in box represents the origin of the *GAL3* allele (YJM978: red, BC187: blue)

To quantify the effect of GAL3, we used a variance-partitioning model with additive effects. We assumed that the total variance of each segregant population (*V_P_*) can be separated into several contributions: *V_P_* = *V_G_* + *V_E_* + *V_EG_* + *V_D_* + *V*_1_. We assumed no interactions between gene and environment (V_EG_=0) and no epistatic interactions (*V_I_*=0). Additionally, there is no dominance as we used haploid strains (V_D_=0) and the environmental variability is equal to the measurement noise because the strains are isogenic and are grown in identical environments (*V_E_*=*ε*^2^). Since we know that *GAL3* is a major driver of the decision threshold phenotype, we partitioned *V_G_* into two components: the variance due to the background (*V_BG_*) and the variance due to *GAL3* (*V_GAL3_*). Hence the total variance could be simplified to *V_P_* = *ε^2^* + *V_GAL3_* + *V_BG_* (Fig. 4, S11 Fig.). By definition, in the allele swap segregants (hybrids 2 and 3) *V_GAL3_* = 0.

Based on this variance-partitioning model (Materials and Methods), we can estimate the contribution of the *GAL3* allele by dividing *V_GAL3_* with the sum of *V_GAL3_* and *V_BG_* or the total genetic variance. We can estimate the *V_BG_* by comparing segregants from hybrid 2 and 3 or from the ‘wild-type’ hybrid, which will give us an upper and lower bound of *GAL3* allelic contribution. Using the segregant population from hybrid 2 and 3, the *GAL3* allele contributes 86% of the genetic variance between YJM978 and BC187. Two segregants from hybrid 1 (‘wildtype’ hybrid) have a decision threshold lower than what we would have expected from segregants of hybrid 2 (S11 Fig.). These two strains increase the background variance, which ultimately reduces the effect of *GAL3*. Using the segregant population from hybrid 1, we estimate that *GAL3* explains 67% of the variance between YJM978 and BC187. These two ‘outliers’ could potentially result from a rare combination of alleles between the strains, implying that we undersampled the distribution from hybrid 2. These calculations suggest that *GAL3* could contribute anywhere from 70-90% to the variance of the decision threshold phenotype.

### Polymorphisms within *GAL3*

To further explore how polymorphisms in *GAL3* might contribute to the decision threshold phenotype, we analyzed the sequences of 55 natural isolates of *S. cerevisiae* [32,46,47]. We identified 8 synonymous and 19 nonsynonymous polymorphisms in the coding region of *GAL3*, which represent 26 unique haplotypes (Table S3). The natural isolates that we assayed (Fig. 1) included 21 of these unique haplotypes, where we excluded the haplotypes from the 6 strains that cannot utilize galactose. To determine whether the *GAL3* haplotype is predictive of the decision threshold, we tested for and found a significant association between decision threshold and *GAL3* haplotype (S12 Fig., p-value=0.04 by ANOVA). However, strains that share *GAL3* haplotypes also tend to share population history (i.e. genomic background) and ecological niche. In particular, YPS163, YPS606, YSP218, and T7 were all isolated from North American oak trees and make up the North American lineage; S288C and FL100 are both mosaic lab strains; YJM978, YJM981, and YJM975, are clinical isolates in the Wine/European lineage. Due to the correlation between phylogeny and *GAL3* haplotype, follow-up investigations using a larger and more diverse set of strains are needed to determine the extent to which decision threshold can be determined solely from the *GAL3* haplotype.

The *GAL3* polymorphisms we observed can in principle affect the expression level, regulation, or function of the protein. Using mutfunc, a database that predicts the consequences of mutations in a protein, we found that 13 of the 19 nonsynonymous SNPs are predicted to affect protein function (Table S4). This includes nonconservative amino-acid substitutions in the Gal3p dimerization interface and the Gal3-Gal80p interface [48]. Gal3p and Gal80p are both homodimers and the Gal3p-Gal80p interaction, which is crucial to the mechanism of GAL pathway activation, is thought to depend on this homodimerization [49,50]. We are less able to predict the impact of promoter variation, but we found 13 SNPs in the promoter (500 bp upstream of the start codon), none of which were in known transcription factor binding sites (Table S3). Furthermore, we did not find a significant association between the *GAL3* promoter haplotype and decision threshold (S12 Fig., p-value=0.98 by ANOVA). Follow-up investigations to characterize the effects of each SNP in *GAL3* will provide mechanistic insight into how the GAL response can be tuned quantitatively by polymorphisms in a single gene.

To determine if *GAL3* or any other genes in the canonical pathway are subject to adaptive evolution, we performed a McDonald-Kreitman analysis [51] using DnaSP [52]. The McDonald-Kreitman test compares intraspecies variation with the divergence between two species. If the ratio of nonsynonymous to synonymous variation between species is equal, there is neutral selection, while any act of natural selection will result in a shift of these two ratios. This test suggests that *GAL3, GAL80*, and *GAL5* are under strong purifying selection (Table S5). Our analysis is consistent with two studies that analyzed polymorphism and divergence data between *S. cerevisiae* and *S. paradoxus*, which suggested that there is strong evidence for purifying selection across the yeast genome [53,54].

### *GAL3* tunes the glucose-galactose diauxic lag

We next asked whether variation in *GAL3* produces selectable variation in phenotype. Diauxic growth is a classical phenotype observed when cells are grown in two sugars [3]. Cells undergo two phases of growth separated by a period with little growth, known as the “diauxic lag”, during which cells induce the genes required to metabolize the second sugar. Previously, our lab has shown that diauxic lag length varies across natural yeast isolates vary, and that *GAL1* transcriptional reporter level before the lag is negatively correlated with lag length [6]. Here, we further show that decision threshold is correlated to *GAL* reporter expression (S13 Fig.), and likely as a result, also negatively correlated with diauxic lag length (Fig. 5A). This suggests that changing *GAL3* alleles will also change the diauxic lag.

**Figure 5.**
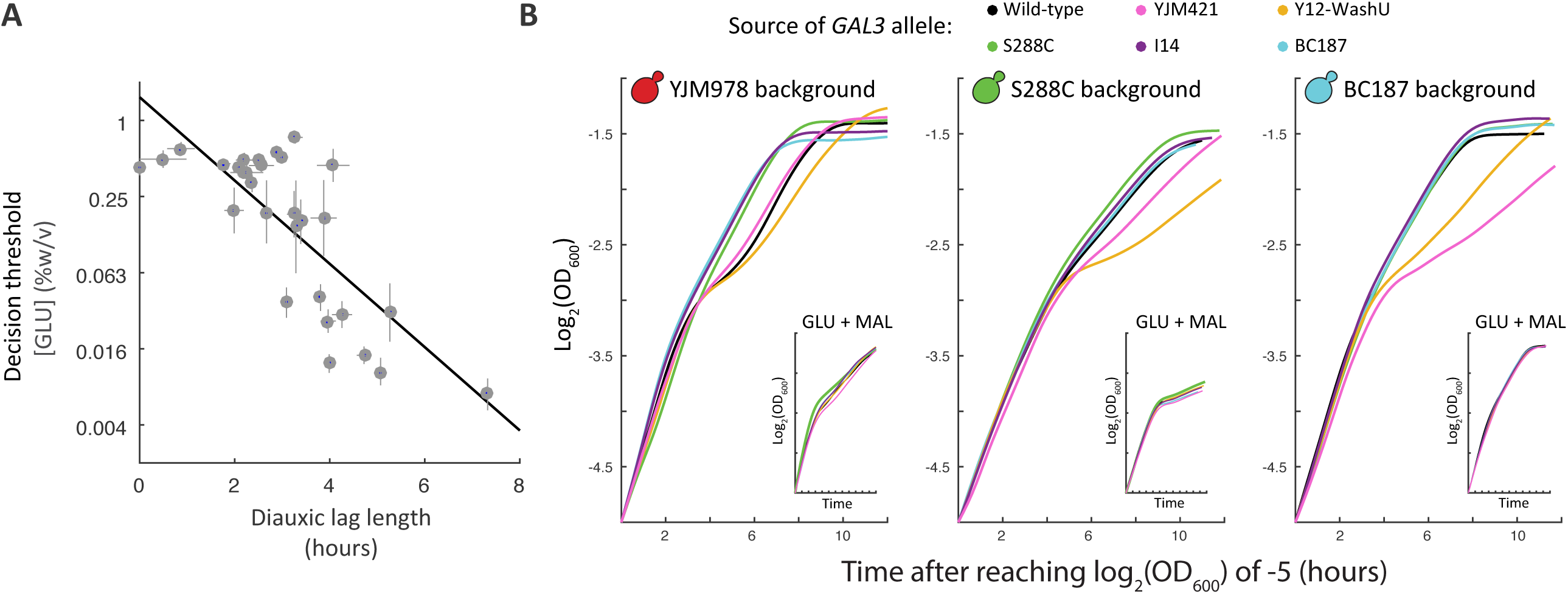
Changing *GAL3* alleles specifically affects the glucose-galactose diauxic lag. (A) Growth curves (OD_600_ versus time) of allele replacement strains in three genetic backgrounds: YJM978 (red), S288C (green), BC187 (blue) of cells grown in a mixture 0.25% glucose and 0.25% galactose. Cultures grew for 6 to 8 hours before entering the diauxic lag. A single replicate is shown (additional replicates are shown in Figure S14). Inset: Growth curves (OD_600_ versus time) of the same strains grown in a mixture of 0.25% glucose and 0.25% maltose. The decision threshold (as measured in Fig. 1) is inversely correlated with the diauxic lag length (as measured in [6]).

To determine if *GAL3* allele affects diauxic lag across our natural isolates, we performed diauxic shift experiments on allele replacement strains representing six *GAL3* alleles (I14, YJM421, Y12-WashU, BC187, and S288c) in three strain backgrounds (YJM978, S288C and BC187) (Fig. 5B). As expected, simply changing the *GAL3* allele in either the YJM978, BC187, or S288C background was sufficient to change the diauxic lag (Fig. 5B, S14 Fig.). Additionally, *GAL3* alleles from short-lag strains S288C, BC187, and I14 (which also have higher decision thresholds) tended to reduce diauxic lag length when introduced into long-lag strain backgrounds YJM978, DBVPG1106, and YJM421, and vice versa. Previously, strains evolved to have an altered diauxic lag in glucose+maltose also had altered lag in glucose+galactose [8]. To determine if the *GAL3* alleles we identified had a specific effect on GAL regulation, we also measured diauxic lag in glucose+maltose. This showed that *GAL3* allele only affects diauxic lag in glucose+galactose and not in glucose+maltose (Fig. 5B, inset).

## Discussion

### Natural genetic variation in the GAL pathway

Genetically and phenotypically diverse natural isolates of yeast have become a powerful system to determine the genetic basis of complex traits. Analyzing natural variation in the well-characterized GAL pathway has the potential to allow us to connect molecular variation, phenotypic variation, and selection. Two recent studies also explored variation in the GAL response across budding yeasts. Peng et al. used combinatorial promoter swaps of GAL regulatory components (*GAL2, GAL3, GAL4, GAL80*) between *S. cerevisiae* and *S. paradoxus* to show that *GAL80* promoter variation was responsible for differences in the GAL response [36]. Roop et al. used a combination of promoter and ORF swaps to show that variation in multiple GAL pathway genes underlies regulatory differences between *S. cerevisiae* and *S. bayanus* [55]. In our study, we used bulk-segregant linkage mapping across diverse *S. cerevisiae* strains to find that most of the variation in GAL regulation is caused by polymorphisms in a single gene, *GAL3*.

Why did each of the three studies identify different genetic loci? A potential explanation is the difference in genetic distance between the strains/species analyzed. We analyzed variation within *S. cerevisiae*, Peng et al. analyzed variation between the closely related species *S. cerevisiae* and *S. paradoxus* [36], and Roop et al. analyzed variation between the more distantly related species *S. cerevisiae* and *S. bayanus* [55]. One hypothesis, based on studies of evolution of development, holds that phenotypic changes on short timescales (i.e. between closely related organisms) are more likely to be caused by nonsynonymous coding-sequence mutations [56,57]. These are favored because of their large phenotypic effects, but come at a cost of increased pleiotropy. On a longer timescale, *cis*-regulatory mutations are enriched, presumably because they are less pleiotropic and allow finely tuned regulation of fitness-enhancing activities [56,57]. Results from Peng et al., Roop et al., and our study are largely inconclusive or weakly inconsistent with pleiotropy being the driving force between the sources of variation. Variation in the GAL response between *S. cerevisiae* and *S. paradoxus* is driven by promoter variation in *GAL80* [58], while variation between *S. cerevisiae* and *S. bayanus* was driven by a combination of promoter and ORF changes [55]. Furthermore, many causative genes were identified between *S. cerevisiae* and *S. bayanus*, while a single gene drove most of the variation within *S. cerevisiae* and between *S. cerevisiae* and *S. paradoxus*. Based on genome-wide expression profiles, there is no evidence that a *GAL80* or *GAL3* variants should be more pleiotropic than simultaneously varying all pathway components [59]. Broader investigations of multiple *Saccharomyces* species will help clarify the relationship between evolutionary distance and the repertoire of mutations.

There are possibilities other than pleiotropy that could cause the difference in genes identified. The different phenotypes assayed in each study could be controlled by different components in the GAL pathway. However, we believe our assays measure highly correlated underlying traits. Peng et al. supplemented their media with mannose to avoid the confounding effects of carbon limitation at low galactose concentrations [36]. The effect of mannose on galactose utilization has not extensively been studied in *S. cerevisiae*, but in other systems mannose can be utilized as a preferred carbon source [60]. Therefore, we expect that the decision threshold in mannose and glucose are likely correlated. Roop et al. compared batch growth in a mixture of glucose and galactose a condition that leads to a diauxic lag in *S. cerevisiae* but not in *S. bayanus*. While natural variation in diauxic growth could have involved many pathways, we showed previously that glucose-galactose diauxic lag is driven by the timing of GAL pathway induction [6]. Here we extended this by showing that diauxic lag is correlated with the decision threshold (Fig. 5A) and primarily modulated by variation in *GAL3* (Fig. 5B). Overall, our work here and recent findings in the literature suggest that all these traits are highly interconnected.

Is the observed variation in the GAL pathway the result of neutral drift or selection? There are three lines of evidence that suggest the GAL pathway is under selection. First, previous analysis has used the QTL cis-regulatory sign test [61] to argue that the GAL pathway has undergone selection between *S. cerevisiae* and *S. bayanus* [55]. Second, we show via the McDonald-Kreitman test that several of the genes in the GAL pathway within *S. cerevisiae* are significantly enriched for nonsynonymous polymorphisms, and therefore likely under adaptive constraint (Table S5). Third, there are at least five genes that affect variability in the GAL response [29,35,36,55]. Given this knowledge we can ask whether the GAL pathway is under selection in a manner similar to the cis-regulatory sign test. Instead of looking for concordant expression changes, we look for enrichment of independent functional mutations in an unexpectedly small subset (i.e. one gene) of multiple possible target genes. Specifically, what is the chance of eight independent alleles of *GAL3* being the main driver of variation in all eight of our crosses given a mutational target size of 5 genes (p-value<1e-6, permutation test). While there are caveats with this method, e.g. what is the true number of potential QTN for each gene, the potential mutational target size is probably much larger than five genes. A recent study of deletion mutants found that upwards of 40% of genes in the yeast genome have the potential to influence the GAL response (Hua et al., under review). Together, we believe these lines of evidence support the hypothesis that the GAL pathway is under selection.

The interplay between selection, pleiotropy, and natural variation is further highlighted by experimental evolution studies in yeast [8] and *E. coli* [62,63]. In mixtures of carbon sources, microbes first consume a preferred nutrient, followed by a “ diauxic lag” where cells must induce the genes necessary to metabolize the second, less preferred nutrient [3]. New et al. found that yeast strains evolved in rapid shifts between glucose and maltose also had a shorter diauxic lag in a mixture of the two sugars [8]. Similarly, *E. coli* passaged in glucose-acetate mixtures evolved into both short-lag and long-lag subpopulations [62,63]. These results show that diauxic lag length is a readily evolvable trait. However, in both previous cases, the evolved phenotypes were due to mutations in global metabolic regulators. For example, New et al. obtained evolved isolates with weakened catabolite repression, via mutations in the glucose-sensing genes *HXK2* and *STD1*, while maltose regulatory genes were unchanged [8]. These mutations are pleiotropic, and thus the evolved strains had shorter diauxic lags in both galactose and maltose. By contrast, we did not find a strong role for general catabolite repression underlying natural variation in GAL regulation, even though the potential mutational target size is large. Instead, our *GAL3* allele replacements specifically tune the glucose-galactose diauxic lag and do not affect the glucose-maltose lag (Fig. 5B, inset). This raises the possibility that in natural environments, where evolution has had longer to act, mutations that perturb global metabolic regulation (as in *STD1* or *HXK2*) may be more detrimental than mutations that tune a particular sugar preference (as in *GAL3*). Hence, as predicted, the frequency of pleiotropic mutations may be an important difference between evolution at short versus long timescales [56,57].

### Genetic complexity of quantitative traits

A number of labs have analyzed phenotypic variation in response to a range of environmental conditions [4,64,65] and delved into the genetic basis of variation in specific traits such as heat tolerance [66], gene/protein expression [67,68], sporulation efficiency [69-71], colony morphology [72-75], sulfur uptake [76], and carbon regulation [36,55]. The vast majority of these studies used growth or expression level [67] as readout. Collectively these studies have yielded insight into the nature of quantitative traits [77]. But, these readouts can potentially miss the complexities of response to fluctuating environments. For example, cells grown in mixtures of glucose and galactose must choose when to induce GAL genes, a property that varies between natural isolates and is distinct from the growth rate on pure glucose and pure galactose (S5 Fig.). Is the decision to induce the GAL pathway similar to other phenotypic traits?

In principle, multiple different pathways and genes can shape natural variation of a trait. A round-robin BSA analysis of MAPK-pathway-mediated stress tolerance in yeast showed that genes both inside and outside the assayed pathway can have large effects on intraspecies phenotypic variation [38]. Previous X-QTL analyses of various traits in yeast have identified a handful of QTLs per trait [38,39,78]. The largest throughput single study found a median of 12 loci for 46 phenotypic traits [79]. While neither previous analysis of the GAL pathway performed a BSA, the number of causative alleles identified through swaps and the total amount of variation explained by these alleles suggests a similar number of genes affect the GAL pathway as other traits [36,55]. Similar to other traits [80], despite the strong correlation between the decision threshold of *GAL3* allele-replacements and their corresponding *GAL3* donor strains, the genetic background still plays a strong role in phenotypic variation (Fig. 3). QTLs outside the GAL pathway might explain why swapping the main regulators of the GAL pathway between *S. cerevisiae* and *S. bayanus* was only able to partially interconvert the phenotypes [55]. Taken together, these results suggest the GAL pathway is similar to other quantitative traits. But, taken alone, our BSA of the GAL pathway suggests that the GAL phenotype is a simpler genetic trait than many of the previously analyzed traits. While other studies have found a small number of QTLs drive the majority of variation in a cross, e.g. sporulation efficiency, when these traits are analyzed in a different cross unique QTLs are often found [70]. Our variation stands out in that there appears to be an allelic series of a single gene, *GAL3*, driving the variation. Similar to the genes whose alleles drive variation in sporulation efficiency, *GAL3* is positioned in a ‘signal transduction bottleneck’ [69]. Unlike sporulation where multiple genes critical for decision making were identified [69], we found variation is driven by a subset, i.e. one, of potential decision making proteins. Taken collectively with the previous analysis of variation in the GAL pathway, an intriguing possibility is this difference might not arise from ‘genetic simplicity’ of the GAL response. Instead, *GAL3* might control the variability in a subset of the phenotype of the GAL response, i.e. the decision threshold, while other members of the GAL pathway might control different aspects of a broader phenotypic response, e.g. diauxic lag. Our future work will directly test this hypothesis.

Genetic interactions, often referred to as epistasis, play a role in many QTL-mapping studies [81,82]. Our study highlights two types of epistasis. First, similar to many other systems [80], the quantitative effect of each *GAL3* allele is influenced by the genetic background. While, the directional effect of the *GAL3* alleles from S288C, BC187, and YJM978 are largely preserved, in the YJM978 and S288C background, the effects of the allele replacements are diminished and the background dominates the resulting phenotype, which is highly compressed (Fig. 3D). This suppression of variation in certain strain backgrounds has been seen in other systems, such as colony morphology [83,84], and can result from the interaction of two or many genes. Second, there appears to be a ‘maximum’ achievable decision threshold of around 1% glucose in 0.25% galactose. Hence, when *GAL3* alleles with decision thresholds above 0.25% glucose in 0.25% galactose are placed into a background with a high decision threshold (e.g. S288C or BC187), the effect of the *GAL3* allele appears to saturate. This behavior is phenomenologically reminiscent of epistatic interactions in peaked fitness landscapes where beneficial mutations have diminishing effects [85,86] or the apparent saturating interaction between gene expression and fitness [87]. Further elucidation of these examples of epistasis in the GAL pathway will likely provide new insights into basic principles of quantitative genetics.

In conclusion, while other genes contribute, the repeated and sizeable role of *GAL3* in this study stands out compared to other QTL analyses in yeast. Using BSA, we identified the galactose sensor *GAL3* as a major driver of this phenotypic variation, accounting for 70-90% of the variation in a single cross. Hence, polymorphisms in a single gene in the canonical GAL pathway are sufficient to create a continuum of natural variation. An intriguing possibility is that in *S. cerevisiae*, variation in *GAL3* may allow strains to vary the diauxic lag in a non-pleiotropic manner. Environments that fluctuate in a ‘predicable’ manner might be expected to select for a pathway architecture that allow strains to evolve on this fluctuating axis [56]. Further analysis of the GAL pathway should help to elucidate the interplay of molecular variation, phenotypic variation, and selection.

## Materials and methods

### Strains and media

Strains were obtained as described in [6]. Strains used in this study can be found in Table S1. All strains from the collection and those assayed in Fig. 1 were homozygous diploids and prototrophic. An initial set of 42 strains were assayed in a gradient of glucose (2% to 0.004% by two-fold dilution) in a background of 0.25% galactose. Strains W303 and YIIC17-E5 were excluded from downstream analysis due to poor growth in our media conditions. Strain 378604X was also excluded due to a high basal expression phenotype that was an outlier in our collection. All experiments were performed in synthetic minimal medium, which contains 1.7g/L Yeast Nitrogen Base (YNB) (BD Difco) and 5g/L ammonium sulfate (EMD), plus D-glucose (EMD), D-galactose (Sigma), or raffinose (Sigma). Cultures were grown in a humidified incubator (Infors Multitron) at 30°C with rotary shaking at 230rpm (tubes and flasks) or 999rpm (600uL cultures in 1mL 96-well plates).

### Flow cytometry assay

GAL induction experiments were performed in a 2-fold dilution series of glucose concentration, from 1% to 0.004% w/v, with constant 0.25% galactose. 2% glucose and 2% galactose conditions were also included with each glucose titration experiment. To assess and control for well-to-well variation, experiments were performed as a co-culture of a “query” strain to be phenotyped and a “reference” strain that was always SLYB93 (natural isolate YJM978 with constitutive mCherry segmentation marker).

To start an experiment, cells were struck onto YPD agar from -80C glycerol stocks, grown to colonies, and then inoculated from colony into YPD liquid and cultured for 16-24 hours. Query and reference strains were then co-innoculated at a 9:1 ratio by volume in a dilution series (1:200 to 1:6400) in S + 2% raffinose medium. The raffinose outgrowths were incubated for 14-16 hours, and then their optical density (OD_600_) was measured on a plate reader (PerkinElmer Envision). One outgrowth culture with OD_600_ closest to 0.1 was selected for each strain, and then washed once in S (0.17% Yeast Nitrogen Base + 0.5% Ammonium Sulfate). Washed cells were diluted 1:200 into glucose + galactose gradients in 96-well plates (500uL cultures in each well) and incubated for 8 hours. Then, cells were processed by washing twice in Tris-EDTA pH 8.0 (TE) and resuspended in TE + 0.1% sodium azide before transferring to a shallow microtiter plate (CELLTREAT) for measurement.

### Calculating the decision threshold (F_50_) metric

Flow cytometry was performed using a Stratedigm S1000EX with A700 automated plate handling system. Data analysis was performed using custom MATLAB scripts, including Flow-Cytometry-Toolkit (https://github.com/springerlab/Flow-Cytometry-Toolkit, https://github.com/springerlab/Induction-Gradient-Toolkit). All experiments were co-cultured with a reference strain and were manually segmented using a fluorescent channel (mCherry or BFP) and side scatter channel (SSC). *GAL1*pr-YFP expression for each segmented population was collected and the induced fraction for each concentration of sugars was computed as shown previously in Escalante et al. [25]. The decision threshold for each glucose titration was calculated from the induced fraction of the ten sugar concentrations. The decision threshold was reported as the glucose concentration were 50% of the cells were induced.

### Filtering reference and query data

To account for well-to-well variability or variability in our glucose titration, each of the query strains were co-cultured with a reference strain, YJM978, containing TDH3pr-mCherry. This constitutive fluorophore was used to segment the query and reference strains. Three filters were used to discard bad samples. 1) The 5% truncated standard deviation was calculated. Samples where the reference strains response was double this truncated deviation from the mean reference response were discarded without analyzing the co-cultured query strain (39 of 480 total experiments) (S2 Fig.). 2) Query strains where the data was of poor quality such that we could not make an accurate calculation of F50, typically for low counts or cultures that did not induce (8 of 441). 3) Query strain values that were over 1.5 standard deviations from the mean of the other replicates, (21 of 433) (S3 Fig.). This 1.5 standard deviation cut-off was determined based on calculating the difference of each sample from the mean and fitting this to a normal distribution assuming outliers (S3 Fig.). All strains were measured at least twice; replicates were performed on different days.

### Estimation of the number of unique GAL phenotypes

To estimate a lower bound for the number of distinct GAL phenotypes, we compared our measurement noise from replicate measurements to the range of variation between strains. By simply dividing the range by the measurement noise or by asking on a pair-wise manner which phenotypes are statistically distinguishable, the number of phenotype is at least five.

### Crossing and generating segregants

To prepare parent strains for crossing and sporulation, diploid natural isolates bearing the *hoΔ::GAL1pr-YFP-hphNT1* reporter cassette were sporulated and random spores were isolated. Mating type was determined by a test cross. We then introduced a constitutive fluorescent marker in tandem with the GAL reporter, to obtain MAT**a**; *hoΔ::GAL1pr-YFP-mTagBFP2-kanMX4* or MATα; *hoΔ::GAL1pr-YFP-mCherry-natMX4* parent strains. To the MAT**a** parent we also introduced a pRS413-derived plasmid bearing *STE2pr-AUR1-C* and *hphNT1*. This plasmid is maintained by hygromycin selection but also allows selection for MAT**a** cells by Aureobasidin A [88]. This plasmid design is inspired by a similar mating-type selection plasmid used in a recent study [38].

To generate segregant pools, we prepared a diploid hybrid and sporulated it as follows. We crossed a parent with *BFP-kanMX* with the mating type selection plasmid to a parent with *mCherry-natMX4* and isolated a G418^R^Nat^R^Hyg^R^ diploid hybrid with the plasmid. We sporulated the hybrid by culturing it to saturation in YPD, diluting 1:10 in YP+2% potassium acetate and incubating at 30C for 8 hours. Cell were then washed and resuspended in 2% potassium acetate and incubated at 30C until >20% of cells were tetrads, or about 72 hours. We incubated ∼5×10^6^ tetrads in 100uL water with 50U of zymolyase 100T (Zymo Research) for 5 hours at 30C, and then resuspended tetrads in 1mL of 1.5% NP-40 and sonicated for 10 seconds at power setting 3 on a probe sonicator (Fisher Scientific Model 550).

To reduce the size of recombination blocks and improve the resolution of linkage mapping [89], we then performed the following “intercross” protocol 4 times: 1) Spores were isolated using the Sony SH800 Cell Sorter selecting for 4×10^6^ BFP+ or mCherry+ (but not +/+ or -/-). 2) The sorted cells were grown into 100uL YPD + 40ug/mL tetracycline. 3) Cells were incubated for 16 hours at 30C without shaking. 4) 5mL of YPD + 200ug/mL G418 + 100ug/mL ClonNat + 200ug/mL Hygromycin B was added and cells were incubated for 48 hours at 30C with shaking. 5) Cultures were sporulated and spores were isolated by zymolyase treatment and sonication as described above. Steps 1-5 were repeated 4 times, resulting in a sonicated suspension of spores that had undergone 5 generations of meiosis since the parents. These spores were resuspended in YPD + 0.5ug/mL AbA and incubated at 30C for 16 hours to select for MAT**a** haploids. This haploid culture was split to create a frozen glycerol stock, and was used as the inoculum for phenotypic isolation by FACS (as described above).

### Sorting-based bulk-segregant analysis

To sort segregant pools for bulk genotyping, the intercrossed MAT**a**-selected segregants were inoculated from a saturated YPD culture into S + 2% raffinose + AbA at dilutions of 1:200, 1:400, 1:800, and 1:1600, and incubated at 30C for 16-24 hours. The outgrowth culture with OD_600_ closest to 0.1 was selected for each strain, washed once in S, and diluted 1:200 into S + 0.25% glucose + 0.25% galactose + AbA. The glucose-galactose culture was incubated at 30C for 8 hours, and then a Sony SH800 sorter was used to isolate pools of 30,000 cells with the 5% lowest (“OFF”) and highest (“ON”) YFP expression, among cells whose Back Scatter (BSC) signal was between 10^5^ and 3×10^5^. This BSC gate was used to minimize the effects of cell size on expression level as cell with similar BSC have similar cell size. The sorted cells were resuspended in YPD + AbA and incubated at 30C until saturation, about 16-24 hours. An aliquot of this culture was saved for -80C glycerol stocks, and another was used to prepare sequencing libraries.

To sequence the segregant pools, genomic DNA was extracted from 0.5mL of saturated YPD culture of each segregant pool using the PureLink Pro 96 kit (Thermo Fisher K182104A). From these genomic preps, sequencing libraries were made using Nextera reagents (Illumina FC-121-1030) following a low-volume protocol [90]. The input DNA concentration was adjusted so that resulting libraries had mean fragment sizes of 200-300bp as measured on a BioAnalyzer. Libraries were multiplexed and sequenced in an Illumina NextSeq flow cell.

### Genome sequences of round-robin parents

Non-S288C parental genomes for the bulk segregant analysis were obtained from the literature: I14 from [38]; BC187, YJM978, DBVPG1106, and Y12 from [91]; YPS606 from [92]. We sequenced our parent strains at ∼1x depth and verified their SNP patterns against these datasets. We initially obtained an unpublished sequence for YJM421 from the NCBI Sequencing Read Archive (accessions SRR097627, SRR096491), but this did not match our strain (it appeared similar to YJM326 instead). A RAD-seq SNP profile of YJM421 [37] partially matched our YJM421, but the RAD-seq data displayed heterozygosity. Because we crossed our YJM421 strain to both I14 and DBVPG1106, for which we have high-quality genomes, we could do the linkage mapping given only one parental genome. However, we confirmed that the YJM421 parent used for both crosses were the same strain, by looking at SNPs in the segregant pools of the two crosses that did not match the other parent. Our current hypothesis is that the YJM421 isolate we obtained from the Fay lab (and which was genotyped by RAD-seq in Cromie et al. [37]) was a heterozygous diploid, a haploid spore of which we used as the parent in our round robin cross.

### Linkage mapping and loci detection

To perform linkage analysis, we aligned raw reads for parent strains (from the literature) and segregant pools (from our experiments) to the sacCer3 (S288C) reference genome using `BWA-MEM` on the Harvard Medical School Orchestra cluster (http://rc.hms.harvard.edu, see Orchestra High Performance Compute Cluster note below). We identified SNPs between cross parents and determined allele counts at each SNP in segregant pools using `samtools mpileup and bcftools call -c`. Using custom MATLAB code, we removed SNPs where read depth was less than 2 or higher than 1000 to avoid alignment artifacts. After filtering, average sequencing depth per pool ranged from 25x to 71x, with a median of 48x.

To calculate LOD scores for allele frequency differences between OFF and ON pools, we input filtered allele counts to the `mp_inference.py` script (MULTIPOOL Version 0.10.2; [41]) with the options `-m contrast -r 100 -c 2200 -n 1000`, following previous practice [38]. A value of n=1000 likely underestimates our segregant pool size and will lead to conservative LOD estimates. An exception to this is the I14xYJM421 cross, which displayed unusually low spore viability (∼20%), possibly due to a Dobzhansky-Muller incompatibility [93]. Thus we used n=200 for this cross.

We defined significant loci as LOD peaks where LOD > 10 (Fig. 2B). Previous bulk segregant analyses using MULTIPOOL used a less stringent cutoff of LOD > 5 [38,39]. This corresponded to a false discovery rate of 5% in one study [39], but led to a much higher number of unreplicated locus calls in another study [38]. Given that our segregant pools underwent multiple rounds of meiosis (and potentially diversity-reducing selection), we chose to use the more conservative LOD > 10. The choice of LOD does not affect our main conclusions about *GAL3*; even the lowest LOD for the chrIV:460 locus (in YJM978 × Y12) is 24 and thus highly significant (S2 Table). Besides this locus, other moderately significant loci may still be biologically relevant, and so we provide a list of LOD peaks and their corresponding support intervals at LOD > 5 (S2 Table). We clustered these peaks as a single locus if they occur within 20kb of each other from different crosses (S8 Fig., S2 Table).

### CRISPR/Cas9 allele replacement

Allele replacement strains were constructed by knocking out *GAL3* (-890bp from start to +911bp from the stop) with KANMX4 followed by CRISPR/Cas9-mediated markerless integration of the heterologous allele. Initially, strains were prepared by introducing Cas9 on a CEN/ARS plasmid (SLVF11); this plasmid is derived from a previous one [94], but the auxotrophic *URA3* marker was replaced with *AUR1-C* to allow Aureobasidin A selection on prototrophic natural isolates. Then, a donor DNA, a guide RNA insert, and a guide RNA backbone were simultaneously transformed into the strain [45]. The donor DNA contained the new allele (PCR amplified from the desired natural isolate genome), its flanking sequences, and an additional 40bp of homology to target it to the correct genomic locus. The guide RNA insert was a linear DNA containing a SNR52 promoter driving a guide RNA gene containing a 20bp CRISPR/Cas recognition sequence linked to a crRNA scaffold sequence, plus 40bp of flanking homology on both ends to a guide RNA backbone. The guide RNA backbone was a 2u plasmid containing natMX4 (pRS420). This was linearized by NotI + XhoI digestion before transformation. Allele re-integration transformations were plated on cloNAT to select for in vivo assembly of the guide RNA into a maintainable plasmid, and Aureobasidin A to select for presence of Cas9. Successful re-integration was verified by colony PCR and Sanger sequencing was performed on a subset of strains and on all donor DNAs to verify the sequence of allelic variants.

### Determining *GAL3* allelic effect by analyzing segregant variance

To estimate the effect of *GAL3* allele on decision threshold, we performed a variance partitioning analysis on decision thresholds of segregants from each of 3 hybrids (Fig. 4). Two heterozygous hybrids with homozygous *GAL3* alleles were constructed by mating CRISPR/Cas9 generated allele replacement strains (YJM978 GAL3^BC187^ or BC187 GAL3^YJM978^) to either BC187 or YJM978 wildtype haploids. A “wildtype” hybrid heterozygous at all loci (BC187 × YM978) was also analyzed. These 3 hybrids were sporulated as described above, and the resulting segregants phenotyped for decision threshold in duplicate.

We assumed a model *V_P_* = *V_GAL3_* + *V_BG_* + *ε*^2^, where phenotypic variance *V_P_* is a sum of contributions from the variance due to *GAL3V_GAL3_*, the variance due to strain background *V_BG_*, and measurement error *ε*^2^. We estimated measurement error by assuming a Gaussian form 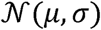 and fitting it to the differences between replicate measurements across all segregants. The variance in inter-replicate differences should be twice the measurement variance, and thus 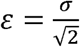. To filter out poor-quality data, we removed segregants where half the inter-replicate difference was greater than 1.5 (S3 Fig.). We calculated the mean of each allele population (*μ_a_* or μ_−a_), where the two allelic variants of *GAL3* are denoted by *a* and −a. To estimate the effect of the *GAL3* allele *E_GAL3_*, we divided the difference of the mean of the two populations by 2. The variance due to *GAL3* is the square of *E_GAL3_*.

Finally, the phenotypic variance of a segregant population (*V_P_*) is composed of the measurement noise (ε^2^) and the genotypic variance (*V_G_*). *V_P_* was calculated for the YJM978 × BC187 segregants and for both of the hybrid conversion segregants. Since *GAL3* is a major driver of the decision threshold phenotype, *V_G_* was partitioned into two components: the contribution to variance of the background (*V_BG_*) and the contribution to variance of *GAL3* (*V_GAL3_*). The background variance was estimated by subtracting *ε*^2^ and *GAL3* variance from the variance of the segregant population. The *GAL3* contribution 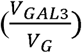 was reported as the ratio of the variance in *GAL3* and the genotypic variance (*V_G_*).

### Growth curves and diauxic lag time metric

Growth curves were obtained as described in Wang et al. [6]. In short, growth curves were obtained by manually measuring the absorbance at 600 nm (OD_600_) on a plate reader (PerkinElmer EnVision) for each plate approximately every 15 min for up to 20 h in a room maintained at 30°C and 75% humidity. Strains to be assayed were pinned into 500 μl of liquid YPD and incubated for 16 h, then diluted 1:200 into 500 μl of synthetic minimal medium + 0.5% glucose and grown for 6-8 h, and finally diluted 1:150 into synthetic minimal medium + 0.25% glucose + 0.25% galactose or synthetic minimal medium + 0.25% glucose + 0.25% maltose for growth curve measurements. The final inoculation was performed into two different plates (with 2 replicates per plate); these replicate growth curves were nearly indistinguishable for all strains. Analysis of growth curve data was performed in MATLAB using custom-written code [6].

To obtain growth rates in glucose or galactose, additional growth curves were performed as above, except the final culture medium contained 0.5% glucose alone or 0.5% galactose alone. The exponential growth rate was extracted from these data as the mean growth rate between when OD_600_ = 2^−6^ and OD_600_ = 2^−4^ (or, equivalently, when culture density was approximately 1/16 and 1/4 of saturation, respectively).

### Bioinformatic analysis

Sequences for the SGRP strains were downloaded from SGRP website. Sequences for the strains in the Liti library [95] were downloaded from https://www.sanger.ac.uk/research/projects/genomeinformatics/sgrp.html. For the remaining strains with multiple distinct isolates reporter in the literature, a single genetic distance that matched the strain in our collection was selected. Using these sequencing databases, we extracted the *GAL3* region and aligned sequences using MUSCLE (Table S3, S12 Fig.). Based on the identified SNPs, we used mutfunc (http://mutfunc.com/) to predict the consequences of nonsynonymous SNPs in the *GAL3* variants (Table S4). These sequences were used for the McDonald Kreitman analysis using DnaSP [52] (Table S5). A neighbor-joining phylogenetic tree was generated using the seqneighjoin function on MATLAB (S7 Fig.) and genetic distances [37].

### Orchestra High Performance Compute Cluster

Portions of this research were conducted on the Orchestra High Performance Compute Cluster at Harvard Medical School. This NIH supported shared facility consists of thousands of processing cores and terabytes of associated storage and is partially provided through grant NCRR 1S10RR028832-01. See http://rc.hms.harvard.edu for more information.

## Acknowledgments

Yarden Katz and Sarah Boswell for critical reading of the manuscript; Andrew Murray and Chiara Ricci-Tam for helpful discussions; and Shervin Javadi and Stratedigm for flow cytometry assistance.

## Supporting information captions

## Supplementary Figures

**Text S1. All supporting figures and captions.**

Contains all the supporting figures and captions in one document.

**Table S1. List of strains used in this study.**

This table lists the strains that were used in this study, Springer Lab ID (all strains used in this study are contained on plate SLL16), genetic background, *GAL3* allele source, genotype, and ecological niche and lineage is listed for diploid strains used in Fig. 1.

**Table S2. List of significant loci and associated genes at LOD > 5.**

This table lists genomic regions for which peak LOD > 5 in the bulk segregant analysis. 2-LOD support intervals are shown for each peak in each cross, as well as averaged support intervals that combine information from “clusters” of peaks within 20kb of each other from different crosses. A subset of genes with sacCer3 (SGD R64-1-1) annotations in the support intervals for each locus are shown.

**Table S3. SNPs found in promoter and coding region of *GAL3* in 55 natural isolates of *S. cerevisiae*.**

This table lists SNPs found in 55 natural isolates of *S. cerevisiae* in the promoter (500bp upstream of start codon) and coding region of *GAL3*. Position on ChrIV is indicated at the top. All genetic variants were compared to the lab strain S288C. 26 different haplotypes are represented across all of the strains.

**Table S4. Predicted structural variants of *GAL3* using mutfunc.**

This table lists the predicted consequences of structural variants found in Gal3p using mutfunc software. Strains that contain the indicated variant are listed in the final column. Mutfunc predicted that 13 out of the 19 nonsynomous mutations have a structural consequence.

**Table S5. McDonald Kreitman analysis of GAL pathway genes.**

This table lists the synonymous and nonsynonymous polymorphisms within *S. cerevisiae* (P_S_/P_N_) compared to synonymous and nonsynonymous between *S. paradoxus* (D_S_/D_N_). SNP counting, pValues, and neutrality indices were analyzed with DnaSP [52].

**S1. Phylogenetic tree of *S. cerevisiae* used in this study.**

Phylogenetic tree of common natural isolates of *S. cerevisiae* constructed based on sequencing data from Cromie et al. 2013 [37]. Strains highlighted in red were used in this study, while strains in black were not.

**S2. Quality control for query strains based on the reference strain.**

Each experiment contained a reference strain. The decision threshold of the reference strain was roughly normally distributed, with a long tail. Based on technical measurements, the tails are due to unintended variation in the assay, e.g. cells grown at too high of an OD, as opposed to biological variation. To eliminate this variation, we truncated the 5% highest and lowest values (red dashed lines). The standard deviation of the remaining, roughly normal, distribution was calculated and used to eliminate samples.

**S3. Determining a cut-off for query outliers.**

(A) Remaining strains were plotted, replicate #1 vs. replicate #2-n, where n is the total number of replicates. A total of 68 strains out of 480 experiments were removed in our quality control. (B) The absolute value of the difference between each distinct measurement of a sample and the mean of all other replicate for that sample is plotted (blue). The same technique was used on simulated derived from a normal distribution of standard deviation 0.75 (red). Based on this a 1.5 standard deviation was chosen to eliminate samples that were likely due to some unintended source of bias.

**S4. Steady-state expression of *GAL1pr-YFP* from a panel of natural isolates in mixtures of glucose and galactose.**

Representative YFP induction profiles of the diploid natural isolates assayed in Fig. 1. Cells were grown for 8 hours, a time previously determined to be sufficient for expression to reach steady-state [25], in a titration from 1% to 0% glucose (two-fold dilution series) in constant background 0.25% galactose. Flow cytometry profiles are plotted for each glucose concentration. Each panel contains 10 distinct glucose and galactose concentrations and 2% pure glucose or galactose. The color density represents the probability density function across of cells for different fluorescent intensity levels. Strains are ordered by increasing decision threshold.

**S5. Growth rate in 0.5% glucose or 0.5% galactose is not strongly correlated with decision threshold.**

Cells were grown in medium containing 0.5% glucose or 0.5% galactose and the OD600 was measured every 15 minutes by plate reader (Materials and methods). The growth rate was then calculated for each strain and condition (Materials and methods). The growth rate in glucose (blue) or galactose (green) of natural isolates is plotted versus the decision threshold (from Fig. 1). Error represents S.E.M. of three replicates for growth rate and at least two replicates for decision threshold. The line is a linear least squared fit.

**S6. Correlation between genetic distance and phenotypic distance for decision threshold and traits from literature.**

Genetic distance [37] and phenotypic distance for a number of traits [4] had been previously measured and determined to be weakly correlated [4]. The histogram of correlation between genetic and phenotypic distance is plotted. The correlation between genetic distance and decision threshold is denoted with the red arrow.

**S7. Relationship of decision threshold with phylogeny and ecological niche.**

Phylogenetic tree was constructed based on the Cromie et al. distance matrix (Materials and methods) with the bar plot indicating decision threshold (from Fig. 1). Color of bars indicate the ecological niche of strain.

**S8. Significance and effect size of detected loci.**

(A) Allele frequency of the ON parent (BC187) in the YJM978xBC187 cross across a region of chromosome IV spanning the chrIV:460Kb locus. The difference in allele frequency between ON and OFF pools at the locus can be used as a proxy for its effect size on the GAL induction phenotype. (B) Scatterplot of significance (LOD score) versus effect size (allele frequency difference) for all 49 LOD peaks where LOD > 5. Significant LOD peaks from different crosses were “clustered” into a single locus if they lay within 20kb of each other. Dots representing LOD peaks are colored by clustered locus.

**S9. Representative YFP induction profiles of *GAL3* allele replacements.**

Homologous *GAL3* allele replacement strains were assayed in a gradient of glucose in a background of 0.25% galactose (Fig. 3A-C). The alleles were assayed in three backgrounds (A) YJM978, (B) BC187, and (C) S288C. (D) The parental strain is shown for comparison.

**S10. Hemizygous hybrids YFP density plots.**

Homologous *GAL3* allele replacement strains were assayed in a gradient of glucose in a background of 0.25% galactose (Fig. 3D). Three different alleles (A) YJM978, (B) BC187, and (C) S288C were assayed in seven genetic backgrounds.

**S11. Phenotypic variation of hybrid (and hybrid conversion) segregants.**

(A) Plot of the decision threshold for replicate 1 and replicate 2 from each segregant assayed. Inset: probability density function of the difference of replicate 1 and replicate 2. The variance from this distribution was used to determine the measurement error. (B) Decision threshold of segregants produced from hybrid conversion (Error represents range of the two segregants).

**S12. Decision threshold versus haplotype of promoter or ORF region.**

Decision threshold plotted versus unique haplotypes of the (A) promoter or (B) ORF region. Haplotype clusters that contain at least two strains are shown on the left side of the graph. The blue line is a guide to show how strains cluster.

**S13. Scatter plot of decision threshold versus *GAL1*-YFP steady state expression [6].**

Scatter plot of steady state *GAL1* expression levels versus decision threshold of a subset of strains from Fig. 1. We previously showed that *GAL1* expression levels before the diauxic lag are inversely correlated with the diauxic lag length [6]. We extend that show that the decision threshold is correlated to these *GAL1* expression levels.

**S14. Growth curves of *GAL3* allele replacement strains.**

Replicate data of growth curves of *GAL3* allele replacement strains in the YJM978, BC187, and S288C background in glucose+galactose (top) or glucose+maltose (bottom). Wild-type growth curves are shown for each background strain in black. Each color represents a different color *GAL3* donor allele. Time is shown relative to Log_2_(OD_600_) reaching -5.

## References

1. Görke B, Stülke J. Carbon catabolite repression in bacteria: many ways to make the most out of nutrients. Nat Rev Micro. 2008;6: 613–624. doi:10.1038/nrmicro1932

2. Gancedo JM. Yeast carbon catabolite repression. Microbiology and Molecular Biology Reviews. 1998.

3. Jacob F, Monod J. Genetic regulatory mechanisms in the synthesis of proteins. Journal of Molecular Biology. Academic Press Inc. (London) Ltd; 1961;3: 318–356. doi:10.1016/S0022-2836(61)80072-7

4. Warringer J, Zörgö E, Cubillos FA, Zia A, Gjuvsland A, Simpson JT, et al. Trait Variation in Yeast Is Defined by Population History. Kruglyak L, editor. PLoS Genet. 2011;7: e1002111–15. doi:10.1371/journal.pgen.1002111

5. Dekel E, Alon U. Optimality and evolutionary tuning of the expression level of a protein. Nature. 2005;436: 588–592. doi:10.1038/nature03842

6. Wang J, Atolia E, Hua B, Savir Y, Escalante-Chong R, Springer M. Natural Variation in Preparation for Nutrient Depletion Reveals a Cost–Benefit Tradeoff. Siegal ML, editor. PLoS Biol. 2015;13: e1002041–31. doi:10.1371/journal.pbio.1002041

7. Lambert G, Kussell E. Memory and Fitness Optimization of Bacteria under Fluctuating Environments. Matic I, editor. PLoS Genet. 2014;10: e1004556–10. doi:10.1371/journal.pgen.1004556

8. New AM, Cerulus B, Govers SK, Perez-Samper G, Zhu B, Boogmans S, et al. Different Levels of Catabolite Repression Optimize Growth in Stable and Variable Environments. Doebeli M, editor. PLoS Biol. 2014;12: e1001764–22. doi:10.1371/journal.pbio.1001764

9. Venturelli OS, Zuleta I, Murray RM, El-Samad H. Population Diversification in a Yeast Metabolic Program Promotes Anticipation of Environmental Shifts. Siegal ML, editor. PLoS Biol. 2015;13: e1002042–24. doi:10.1371/journal.pbio.1002042

10. Sorrells TR, Booth LN, Tuch BB, Johnson AD. Intersecting transcription networks constrain gene regulatory evolution. Nature. 2015;523: 361–365. doi:10.1038/nature14613

11. Borneman AR, Chambers PJ, Pretorius IS. Yeast systems biology: modelling the winemaker’s art. Trends Biotechnol. 2007;25: 349–355. doi:10.1016/j.tibtech.2007.05.006

12. Tuch BB, Li H, Johnson AD. Evolution of Eukaryotic Transcription Circuits. Science. American Association for the Advancement of Science; 2008;319: 1797–1799. doi:10.1126/science.1152398

13. Galgoczy DJ, Cassidy-Stone A, Llinas M, O’Rourke SM, Herskowitz I, DeRisi JL, et al. Genomic dissection of the cell-type-specification circuit in Saccharomyces cerevisiae. Proceedings of the National Academy of Sciences. National Acad Sciences; 2004;101: 18069–18074. doi:10.1073/pnas.0407611102

14. Odom DT, Dowell RD, Jacobsen ES, Gordon W, Danford TW, MacIsaac KD, et al. Tissue-specific transcriptional regulation has diverged significantly between human and mouse. Nat Genet. Nature Publishing Group; 2007;39: 730–732. doi:10.1038/ng2047

15. Bradley RK, Li X-Y, Trapnell C, Davidson S, Pachter L, Chu HC, et al. Binding Site Turnover Produces Pervasive Quantitative Changes in Transcription Factor Binding between Closely Related Drosophila Species. Wray GA, editor. PLoS Biol. Public Library of Science; 2010;8: e1000343. doi:10.1371/journal.pbio.1000343

16. Dalal CK, Zuleta IA, Mitchell KF, Andes DR. Transcriptional rewiring over evolutionary timescales changes quantitative and qualitative properties of gene expression. eLife. 2016. doi:10.7554/eLife.18981.001

17. Tsong AE, Miller MG, Raisner RM, Johnson AD. Evolution of a Combinatorial Transcriptional Circuit. Cell. 2003;115: 389–399. doi:10.1016/S0092-8674(03)00885-7

18. Tsong AE, Tuch BB, Li H, Johnson AD. Evolution of alternative transcriptional circuits with identical logic. Nature. Nature Publishing Group; 2006;443: 415–420. doi:10.1038/nature05099

19. Lavoie H, Hogues H, Whiteway M. Rearrangements of the transcriptional regulatory networks of metabolic pathways in fungi. Current Opinion in Microbiology. 2009;12: 655–663. doi:10.1016/j.mib.2009.09.015

20. Dalal CK, Zuleta IA, Mitchell KF, Andes DR. Transcriptional rewiring over evolutionary timescales changes quantitative and qualitative properties of gene expression. eLife. 2016. doi:10.7554/eLife.18981.001

21. Hittinger CT, Carroll SB. Gene duplication and the adaptive evolution of a classic genetic switch. Nature. 2007;449: 677–681. doi:10.1038/nature06151

22. Nehlin JO, Carlberg M, Ronne H. Control of yeast GAL genes by MIG1 repressor: a transcriptional cascade in the glucose response. The EMBO Journal. European Molecular Biology Organization; 1991;10: 3373–3377.

23. Johnston M, Flick JS, Pexton T. Multiple mechanisms provide rapid and stringent glucose repression of GAL gene expression in Saccharomyces cerevisiae. Molecular and Cellular Biology. American Society for Microbiology; 1994;14: 3834–3841. doi:10.1128/MCB.14.6.3834

24. Bennett MR, Pang WL, Ostroff NA, Baumgartner BL, Nayak S, Tsimring LS, et al. Metabolic gene regulation in a dynamically changing environment. Nature. 2008;454: 1119–1122. doi:10.1038/nature07211

25. Escalante-Chong R, Savir Y, Carroll SM, Ingraham JB, Wang J, Marx CJ, et al. Galactose metabolic genes in yeast respond to a ratio of galactose and glucose. Proceedings of the National Academy of Sciences. 2015;112: 1636–1641. doi:10.1073/pnas.1418058112

26. Bhat PJ. Galactose Regulon of Yeast. Berlin, Heidelberg: Springer Berlin Heidelberg; 2008. doi:10.1007/978-3-540-74015-5

27. Scott A, Timson DJ. Characterization of the Saccharomyces cerevisiae galactose mutarotase/UDP-galactose 4-epimerase protein, Gal10p. FEMS Yeast Research. The Oxford University Press; 2007;7: 366–371. doi:10.1111/j.1567- 1364.2006.00204.x

28. Lohr D, Venkov P, Zlatanova J. Transcriptional regulation in the yeast GAL gene family: a complex genetic network. FASEB J. Federation of American Societies for Experimental Biology; 1995;9: 777–787. doi:10.1096/fj.1530-6860

29. Acar M. A General Mechanism for Network-Dosage Compensation in Gene Circuits. Science. 2010;329: 1656–1660. doi:10.1126/science.1190544

30. Acar M. Enhancement of cellular memory by reducing stochastic transitions. Nature. 2005;435: 228–232. doi:10.1038/nature03524

31. Baganz F, Hayes A, Marren D, Gardner DCJ, Oliver SG. Suitability of replacement markers for functional analysis studies inSaccharomyces cerevisiae. Yeast. John Wiley & Sons, Ltd; 1997;13: 1563–1573. doi:10.1002/(SICI)1097-0061(199712)13:16<1563::AID-YEA240>3.3.CO;2-Y

32. Liti G, Carter DM, Moses AM, Warringer J, Parts L, James SA, et al. Population genomics of domestic and wild yeasts. Nature. 2009;458: 337–341. doi:10.1038/nature07743

33. Fay JC, Benavides JA. Evidence for Domesticated and Wild Populations of Saccharomyces cerevisiae. PLoS Genet. 2005;1: e5–6. doi:10.1371/journal.pgen.0010005

34. Verma M, Bhat PJ, Venkatesh KV. Quantitative Analysis of GAL Genetic Switch of Saccharomyces cerevisiae Reveals That Nucleocytoplasmic Shuttling of Gal80p Results in a Highly Sensitive Response to Galactose. Journal of Biological Chemistry. 2003;278: 48764–48769. doi:10.1074/jbc.M303526200

35. Venturelli OS, El-Samad H, Murray RM. Synergistic dual positive feedback loops established by molecular sequestration generate robust bimodal response. Proceedings of the National Academy of Sciences. 2012;109: E3324–E3333. doi:10.1073/pnas.1211902109

36. Peng W, Liu P, Xue Y, Acar M. Evolution of gene network activity by tuning the strength of negative-feedback regulation. Nature Communications. Nature Publishing Group; 1AD;6: 1–9. doi:10.1038/ncomms7226

37. Cromie GA, Hyma KE, Ludlow CL, Garmendia-Torres C, Gilbert TL, May P, et al. Genomic Sequence Diversity and Population Structure of Saccharomyces cerevisiae Assessed by RAD-seq. G3&#58; Genes|Genomes|Genetics. 2013;3: 2163–2171. doi:10.1534/g3.113.007492

38. Treusch S, Albert FW, Bloom JS, Kotenko IE, Kruglyak L. Genetic Mapping of MAPK-Mediated Complex Traits Across S. cerevisiae. Copenhaver GP, editor. PLoS Genet. 2015;11: e1004913–16. doi:10.1371/journal.pgen.1004913

39. Albert FW, Treusch S, Shockley AH, Bloom JS, Kruglyak L. Genetics of single-cell protein abundance variation in large yeast populations. Nature. Nature Publishing Group; 2014;506: 494–497. doi:10.1038/nature12904

40. Ehrenreich IM, Torabi N, Jia Y, Kent J, Martis S, Shapiro JA, et al. Dissection of genetically complex traits with extremely large pools of yeast segregants. Nature. Nature Publishing Group; 2010;464: 1039–1042. doi:10.1038/nature08923

41. Edwards MD, Gifford DK. High-resolution genetic mapping with pooled sequencing. BMC Bioinformatics. BioMed Central Ltd; 2012;13: S8. doi:10.1186/1471-2105-13-S6-S8

42. Lavy T, Kumar PR, He H, Joshua-Tor L. The Gal3p transducer of the GAL regulon interacts with the Gal80p repressor in its ligand-induced closed conformation. Genes & Development. 2012;26: 294–303. doi:10.1101/gad.182691.111

43. Lorenz K, Cohen BA. Small- and Large-Effect Quantitative Trait Locus Interactions Underlie Variation in Yeast Sporulation Efficiency. Genetics. 2012;192: 1123–1132. doi:10.1534/genetics.112.143107

44. Duveau F, Metzger B, Gruber JD, Mack K. Mapping small effect mutations in Saccharomyces cerevisiae: impacts of experimental design and mutational properties. G3: Genes| Genomes|…. 2014. doi:10.1534/g3.114.011783/-/DC1

45. Horwitz AA, Walter JM, Schubert MG, Kung SH, Hawkins K, Platt DM, et al. Efficient Multiplexed Integration of Synergistic Alleles and Metabolic Pathways in Yeasts via CRISPR-Cas. Cell Systems. The Authors; 2015;1: 88–96. doi:10.1016/j.cels.2015.02.001

46. Schacherer J, Shapiro JA, Ruderfer DM, Kruglyak L. Comprehensive polymorphism survey elucidates population structure of Saccharomyces cerevisiae. Nature. 2009;458: 342–345. doi:10.1038/nature07670

47. Bergstrom A, Simpson JT, Salinas F, Barre B, Parts L, Zia A, et al. A High-Definition View of Functional Genetic Variation from Natural Yeast Genomes. Molecular Biology and Evolution. 2014;31: 872–888. doi:10.1093/molbev/msu037

48. Lavy T, Kumar PR, He H, Joshua-Tor L. The Gal3p transducer of the GAL regulon interacts with the Gal80p repressor in its ligand-induced closed conformation. Genes & Development. 2012;26: 294–303. doi:10.1101/gad.182691.111

49. Melcher K, Xu HE. Gal80-Gal80 interaction on adjacent Gal4p binding sites is required for complete GAL gene repression. The EMBO Journal. 2001.

50. Egriboz O, Goswami S, Tao X, Dotts K, Schaeffer C, Pilauri V, et al. Self-Association of the Gal4 Inhibitor Protein Gal80 Is Impaired by Gal3: Evidence for a New Mechanism in the GAL Gene Switch. Molecular and Cellular Biology. 2013;33: 3667–3674. doi:10.1128/MCB.00646-12

51. McDonald JH, Kreitman M. Adaptive protein evolution at the Adh locus in Drosophila. Nature. 1991.

52. Rozas J. DNA Sequence Polymorphism Analysis Using DnaSP. In: Mapelli V, editor. Yeast Metabolic Engineering. Totowa, NJ: Humana Press; 2009. pp. 337–350. doi:10.1007/978-1-59745-251-9_17

53. Elyashiv E, Bullaughey K, Sattath S, Rinott Y, Przeworski M, Sella G. Shifts in the intensity of purifying selection: An analysis of genome-wide polymorphism data from two closely related yeast species. Genome Research. 2010;20: 1558–1573. doi:10.1101/gr.108993.110

54. Doniger SW, Kim HS, Swain D, Corcuera D, Williams M, Yang S-P, et al. A Catalog of Neutral and Deleterious Polymorphism in Yeast. Pritchard JK, editor. PLoS Genet. 2008;4: e1000183–15. doi:10.1371/journal.pgen.1000183

55. Roop JI, Chang KC, Brem RB. Polygenic evolution of a sugar specialization trade-off in yeast. Nature. Nature Publishing Group; 2016;530: 336–339. doi:10.1038/nature16938

56. Stern DL, Orgogozo V. THE LOCI OF EVOLUTION: HOW PREDICTABLE IS GENETIC EVOLUTION? Evolution. 2008;62: 2155–2177. doi:10.1111/j.1558-5646.2008.00450.x

57. Stern DL. Is Genetic Evolution Predictable? Science. 2009;323: 741–746. doi:10.1126/science.1158997

58. Peng W, Liu P, Xue Y, Acar M. Evolution of gene network activity by tuning the strength of negative-feedback regulation. Nature Communications. Nature Publishing Group; 1AD;6: 1–9. doi:10.1038/ncomms7226

59. Lashkari DA, DeRisi JL, McCusker JH. Yeast microarrays for genome wide parallel genetic and gene expression analysis. 1997.

60. Monod J. The growth of bacterial cultures. Annual Reviews in Microbiology. 1949.

61. Orr HA. Testing natural selection vs. genetic drift in phenotypic evolution using quantitative trait locus data. Genetics. 1998.

62. Friesen ML, Saxer G, Travisano M, Doebeli M. EXPERIMENTAL EVIDENCE FOR SYMPATRIC ECOLOGICAL DIVERSIFICATION DUE TO FREQUENCY-DEPENDENT COMPETITION IN ESCHERICHIA COLI. Evolution. 2004;58: 245–17. doi:10.1554/03-369

63. Spencer CC, Saxer G. Seasonal resource oscillations maintain diversity in bacterial microcosms. Evolutionary Ecology…. 2007.

64. Smith EN, Kruglyak L. Gene–Environment Interaction in Yeast Gene Expression. Mackay T, editor. PLoS Biol. 2008;6: e83–15. doi:10.1371/journal.pbio.0060083

65. Ehrenreich IM, Torabi N, Jia Y, Kent J, Martis S, Shapiro JA, et al. Dissection of genetically complex traits with extremely large pools of yeast segregants. Nature. Nature Publishing Group; 2010;464: 1039–1042. doi:10.1038/nature08923

66. Steinmetz LM, Sinha H, Richards DR, Spiegelman JI, Oefner PJ, McCusker JH, et al. Dissecting the architecture of a quantitative trait locus in yeast. Nature. 2002;416: 326–330. doi:10.1038/416326a

67. Brem RB. Genetic Dissection of Transcriptional Regulation in Budding Yeast. Science. 2002;296: 752–755. doi:10.1126/science.1069516

68. Yvert G, Brem RB, Whittle J, Akey JM, Foss E, Smith EN, et al. Trans-acting regulatory variation in Saccharomyces cerevisiae and the role of transcription factors. Nat Genet. 2003;35: 57–64. doi:10.1038/ng1222

69. Lorenz K, Cohen BA. Causal Variation in Yeast Sporulation Tends to Reside in a Pathway Bottleneck. Gibson G, editor. PLoS Genet. 2014;10: e1004634–12. doi:10.1371/journal.pgen.1004634

70. Gerke JP, Chen CTL, Cohen BA. Natural Isolates of Saccharomyces cerevisiae Display Complex Genetic Variation in Sporulation Efficiency. Genetics. 2006;174: 985–997. doi:10.1534/genetics.106.058453

71. Gerke J. Genetic interactions between transcription factors cause natural variation in yeast. Science. 2009;323: 495–498. doi:10.1126/science.1166426

72. Ruusuvuori P, Lin J, Scott AC, Tan Z, Sorsa S, Kallio A, et al. Quantitative analysis of colony morphology in yeast. BioTechniques. 2014;56: 1–14. doi:10.2144/000114123

73. Taylor MB, Ehrenreich IM. Genetic Interactions Involving Five or More Genes Contribute to a Complex Trait in Yeast. Fay JC, editor. PLoS Genet. 2014;10: e1004324–8. doi:10.1371/journal.pgen.1004324

74. Voordeckers K, De Maeyer D, van der Zande E, Vinces MD, Meert W, Cloots L, et al. Identification of a complex genetic network underlying Saccharomyces cerevisiaecolony morphology. Molecular Microbiology. 2012;86: 225–239. doi:10.1111/j.1365-2958.2012.08192.x

75. Granek JA, Magwene PM. Environmental and genetic determinants of colony morphology in yeast. PLoS Genet. 2010. doi:10.1371/journal.pgen.1000823.g001

76. Sanchez MR, Miller AW, Liachko I, Sunshine AB, Lynch B, Huang M, et al. Differential paralog divergence modulates genome evolution across yeast species. Fay JC, editor. PLoS Genet. 2017;13: e1006585–27. doi:10.1371/journal.pgen.1006585

77. Fay JC. The molecular basis of phenotypic variation in yeast. Current Opinion in Genetics & Development. Elsevier Ltd; 2013;23: 672–677. doi:10.1016/j.gde.2013.10.005

78. Ehrenreich IM, Bloom J, Torabi N, Wang X, Jia Y, Kruglyak L. Genetic Architecture of Highly Complex Chemical Resistance Traits across Four Yeast Strains. Gasch AP, editor. PLoS Genet. 2012;8: e1002570–9. doi:10.1371/journal.pgen.1002570

79. Bloom JS, Kotenko I, Sadhu MJ, Treusch S, Albert FW, Kruglyak L. Genetic interactions contribute less than additive effects to quantitative trait variation in yeast. Nature Communications. Nature Publishing Group; 2015;6: 1–6. doi:10.1038/ncomms9712

80. Chandler CH, Chari S, Dworkin I. Does your gene need a background check? How genetic background impacts the analysis of mutations, genes, and evolution. Trends in Genetics. Elsevier Ltd; 2013;29: 358–366. doi:10.1016/j.tig.2013.01.009

81. Bloom JS, Ehrenreich IM, Loo WT, Lite T-LV, Kruglyak L. Finding the sources of missing heritability in a yeast cross. Nature. Nature Publishing Group; 2013;: 1–6. doi:10.1038/nature11867

82. Phillips PC. Epistasis — the essential role of gene interactions in the structure and evolution of genetic systems. Nat Rev Genet. 2008;9: 855–867. doi:10.1038/nrg2452

83. Taylor MB, Ehrenreich IM. Genetic Interactions Involving Five or More Genes Contribute to a Complex Trait in Yeast. Fay JC, editor. PLoS Genet. 2014;10: e1004324–8. doi:10.1371/journal.pgen.1004324

84. Lee JT, Taylor MB, Shen A, Ehrenreich IM. Multi-locus Genotypes Underlying Temperature Sensitivity in a Mutationally Induced Trait. Siegal ML, editor. PLoS Genet. 2016;12: e1005929–18. doi:10.1371/journal.pgen.1005929

85. Kryazhimskiy S, Rice DP, Jerison ER, Desai MM. Global epistasis makes adaptation predictable despite sequence-level stochasticity. Science. 2014.

86. Plucain J, Hindré T, Le Gac M, Tenaillon O. Epistasis and allele specificity in the emergence of a stable polymorphism in Escherichia coli.… 2014. doi:10.1126/science.1248688

87. Keren L, Hausser J, Lotan-Pompan M, Slutskin IV, Alisar H, Kaminski S, et al. Massively Parallel Interrogation of the Effects of Gene Expression Levels on Fitness. Cell. Elsevier Inc; 2016;: 1–32. doi:10.1016/j.cell.2016.07.024

88. Hashida-Okado T, Ogawa A, Kato I, Takesako K. Transformation system for prototrophic industrial yeasts using the AUR1gene as a dominant selection marker. FEBS Letters. 1998;425: 117–122. doi:10.1016/S0014-5793(98)00211-7

89. Cubillos FA, Parts L, Salinas F, Bergstrom A, Scovacricchi E, Zia A, et al. High-Resolution Mapping of Complex Traits with a Four-Parent Advanced Intercross Yeast Population. Genetics. 2013;195: 1141–1155. doi:10.1534/genetics.113.155515

90. Baym M, Kryazhimskiy S, Lieberman TD, Chung H, Desai MM, Kishony R. Inexpensive Multiplexed Library Preparation for Megabase-Sized Genomes. Green SJ, editor. PLoS ONE. 2015;10: e0128036–15. doi:10.1371/journal.pone.0128036

91. Bergstrom A, Simpson JT, Salinas F, Barre B, Parts L, Zia A, et al. A High-Definition View of Functional Genetic Variation from Natural Yeast Genomes.Molecular Biology and Evolution. 2014;31: 872–888. doi:10.1093/molbev/msu037

92. Strope PK, Skelly DA, Kozmin SG, Mahadevan G, Stone EA, Magwene PM, et al. The 100-genomes strains, an S. cerevisiaeresource that illuminates its natural phenotypic and genotypic variation and emergence as an opportunistic pathogen. Genome Research. 2015;25: 762–774. doi:10.1101/gr.185538.114

93. Hou J, Friedrich A, Gounot J-S, Schacherer J. Comprehensive survey of condition-specific reproductive isolation reveals genetic incompatibility in yeast. Nature Communications. Nature Publishing Group; 2015;6: 7214. doi:10.1038/ncomms8214

94. DiCarlo JE, Norville JE, Mali P, Rios X, Aach J, Church GM. Genome engineering in Saccharomyces cerevisiae using CRISPR-Cas systems. Nucleic Acids Research. 2013;41: 4336–4343. doi:10.1093/nar/gkt135

95. Liti G, Louis EJ. Advances in Quantitative Trait Analysis in Yeast. Fay JC, editor. PLoS Genet. 2012;8: e1002912–7. doi:10.1371/journal.pgen.1002912

